# Regulation of KHNYN antiviral activity by the extended di-KH domain and nucleo-cytoplasmic trafficking

**DOI:** 10.1101/2021.12.22.473955

**Authors:** María José Lista, Rui Pedro Galão, Mattia Ficarelli, Dorota Kmiec, Harry Wilson, Helena Winstone, Elizabeth R Morris, Hannah E Mischo, Joseph Wanford, Rebecca Youle, Charlotte Odendall, Ian A Taylor, Stuart J D Neil, Chad M Swanson

**Affiliations:** King’s College London, Department of Infectious Diseases, London, United Kingdom; The Francis Crick Institute, Macromolecular Structure Laboratory, London, United Kingdom

## Abstract

The zinc finger antiviral protein (ZAP) restricts a broad range of viruses by binding CpG dinucleotides in viral RNA to target it for degradation and inhibit its translation. KHNYN was recently identified as an antiviral protein required for ZAP to inhibit retroviral replication, though little is known about its functional determinants. KHNYN contains an N-terminal extended di-KH-like domain, a PIN endoribonuclease domain and a C-terminal CUBAN domain that binds NEDD8 and ubiquitin. We show that deletion of the extended di-KH domain reduces its antiviral activity. However, despite its similarity to RNA binding KH domains, the extended di-KH domain in KHNYN does not appear to bind RNA. Mutation of residues in the CUBAN domain that bind NEDD8 increase KHNYN abundance but do not alter its antiviral activity, suggesting that this interaction regulates KHNYN homeostatic turnover. In contrast, a CRM1-dependent nuclear export signal (NES) at the C-terminus of the CUBAN domain is required for antiviral activity. Deletion of this signal retains KHNYN in the nucleus and inhibits its interaction with ZAP. Interestingly, this NES appeared in the KHNYN lineage at a similar time as when ZAP evolved in tetrapods, indicating that these proteins may have co-evolved to restrict viral replication.

**AUTHOR SUMMARY:** Antiviral proteins restrict viral replication in many different ways, including inhibiting viral gene expression. ZAP is an antiviral RNA binding protein that must interact with other cellular proteins to inhibit viral protein synthesis. KHNYN is a ZAP cofactor that is required for it to inhibit retroviral replication. Because little is known about how KHNYN functions in this role, we have analyzed how two of its domains regulate its antiviral activity. We first show that the extended di-KH domain in KHNYN is required for its antiviral activity. While it is related to di-KH domains in RNA binding proteins, it appears to have lost its ability to bind RNA and KHNYN likely acts in the restriction pathway after ZAP binds a target viral RNA. Second, we show that the KHNYN CUBAN domain regulates both its protein abundance and trafficking within the cell. The CUBAN domain contains a nuclear export signal and, when this signal is mutated, KHNYN is sequestered in the nucleus, has substantially reduced antiviral activity and does not interact with ZAP. Overall, we show that the extended di-KH and CUBAN domains in KHNYN are required for it to act as a cofactor for ZAP to restrict viral replication.

## INTRODUCTION

The zinc finger antiviral protein (ZAP, also known as ZC3HAV1 or PARP13) is an RNA binding protein that targets viral RNA containing CpG dinucleotides for degradation and inhibits their translation [1]. Unlike many viral RNA sensors, ZAP binds single stranded RNA instead of double stranded RNA, allowing it to potentially bind cellular mRNAs [2–4]. Furthermore, as very few cellular transcripts are devoid of CpGs, ZAP activity must be highly regulated to prevent it from targeting many cellular mRNAs and causing genome-wide changes in gene expression [1, 5, 6]. ZAP has no intrinsic catalytic activity and has been reported to interact with multiple cofactors to mediate RNA degradation including DDX17, the RNA exosome complex, PARN and TRIM25, though these are not required to inhibit all ZAP-sensitive viruses [7–11]. Therefore, determining how ZAP cofactors contribute to its activity is essential to understand how this antiviral system mediates potent and selective restriction. We recently identified KHNYN, a putative endoribonuclease with no previously known function, as a cofactor required for ZAP to inhibit retroviral replication [12].

The name KHNYN is derived from its original annotation that suggested that it contains a type I K homology (KH) domain and a Nedd4-BP1, YacP-like Nuclease (NYN) domain [13]. The N-terminal KH domain has been reported to be unusual due to a metal chelating domain insertion [13], though this has not been characterized. The NYN domain was originally reported to be similar to the PilT N-terminal (PIN) nuclease fold and a recent classification of the PIN domain-like superfamily reclassified this domain in KHNYN from the NYN group to the proteinaceous RNase P (PRORP) group containing the PRORP enzymes and the ZC3H12 RNase family [13, 14]. All PIN domains have a catalytic core formed from four conserved aspartic acid residues that coordinate a Mg^2+^ ion and we have shown that mutating these conserved aspartic acid residues eliminates KHNYN antiviral activity [12, 14]. In addition to the KH and PIN domains, KHNYN also has a C-terminal cullin-binding domain associating with NEDD8 (CUBAN). This domain binds NEDD8 and ubiquitin, both of which are members of the ubiquitin-like family, and preferentially binds monomeric NEDD8 over ubiquitin [15]. NEDD8 binding mediates an interaction between KHNYN and neddylated cullin–RING E3 ubiquitin ligases. However, the role of the CUBAN domain for KHNYN antiviral activity, or any other function, is not known.

KHNYN has two human paralogs, NYNRIN and N4BP1. NYNRIN evolved from a KHNYN gene duplication in which the RNase H and integrase domains from an endogenous retrovirus replaced the last exon of KHNYN [16]. The function of this protein is unknown. N4BP1 contains domains that are paralogous to the KH domain, PIN domain and CUBAN domains in KHNYN and it is a predominantly nucleolar protein whose expression is induced by type I interferon [13, 17-20]. While the specific functions of N4BP1 are still unclear, it has been shown to inhibit the NF-KB pathway as well as HIV-1 gene expression and the E3 ligase Itch [21–24]. N4BP1 also has a genetic interaction with ZAP in that ZAP is required for N4BP1 antiviral activity [25].

Human immunodeficiency virus type 1 (HIV-1) is a common model system to study the antiviral activity of ZAP and its cofactors because it is highly depleted in CpG dinucleotides, which makes it poorly targeted by ZAP [26–28]. However, when a specific region in HIV-1 *env* is engineered to contain additional CpGs through synonymous mutations, the virus becomes ZAP-sensitive [28–30]. There are two predominant isoforms for ZAP, ZAP-L and ZAP-S [31, 32]. ZAP-L contains a C-terminal S-farnesylation modification that localizes it to the cytoplasmic endomembrane system and it has greater antiviral activity than ZAP-S against some viruses, including CpG-enriched HIV-1 [32–36]. Importantly, KHNYN physically interacts with ZAP and is required for ZAP to restrict retroviruses, including CpG-enriched HIV-1 [12, 29].

In this study, we characterized how the KH domain and CUBAN domain regulate KHNYN antiviral activity. In contrast to its original annotation [13], the KH domain is composed of an extended di-KH domain that appears to be unique to the KHNYN/N4BP1/NYNRIN family. While it is required for full antiviral activity, it likely does not bind RNA, at least in a conventional manner. The CUBAN domain has at least two functions for KHNYN. First, its ability to bind NEDD8 is required for KHNYN homeostatic turnover but not its antiviral activity. Second, the extreme C-terminus of the CUBAN domain contains a CRM1 nuclear export signal (NES) that is required for proper localization of KHNYN to the cytoplasm. This regulates its interaction with ZAP and is essential for antiviral activity. Furthermore, the NES in KHNYN appears to have co-evolved with ZAP as two of the five key residues in the NES emerged at the same time as ZAP evolved in tetrapods.

## RESULTS

### The extended di-KH domain in KHNYN is required for its antiviral activity

To characterize the domains in KHNYN, we used the Phyre2 protein structure prediction tool [37]. This identified three major domains: an N-terminal di-KH domain (residues 10-195), a PIN domain (residues 436-595) and the C-terminal CUBAN domain (residues 627-678) as well as a long inter-domain region (residues 196-435) that is predicted to be largely disordered with low hydrophobicity (Figure 1A). A similar model for this protein was predicted by AlphaFold [38].

**Figure 1.**
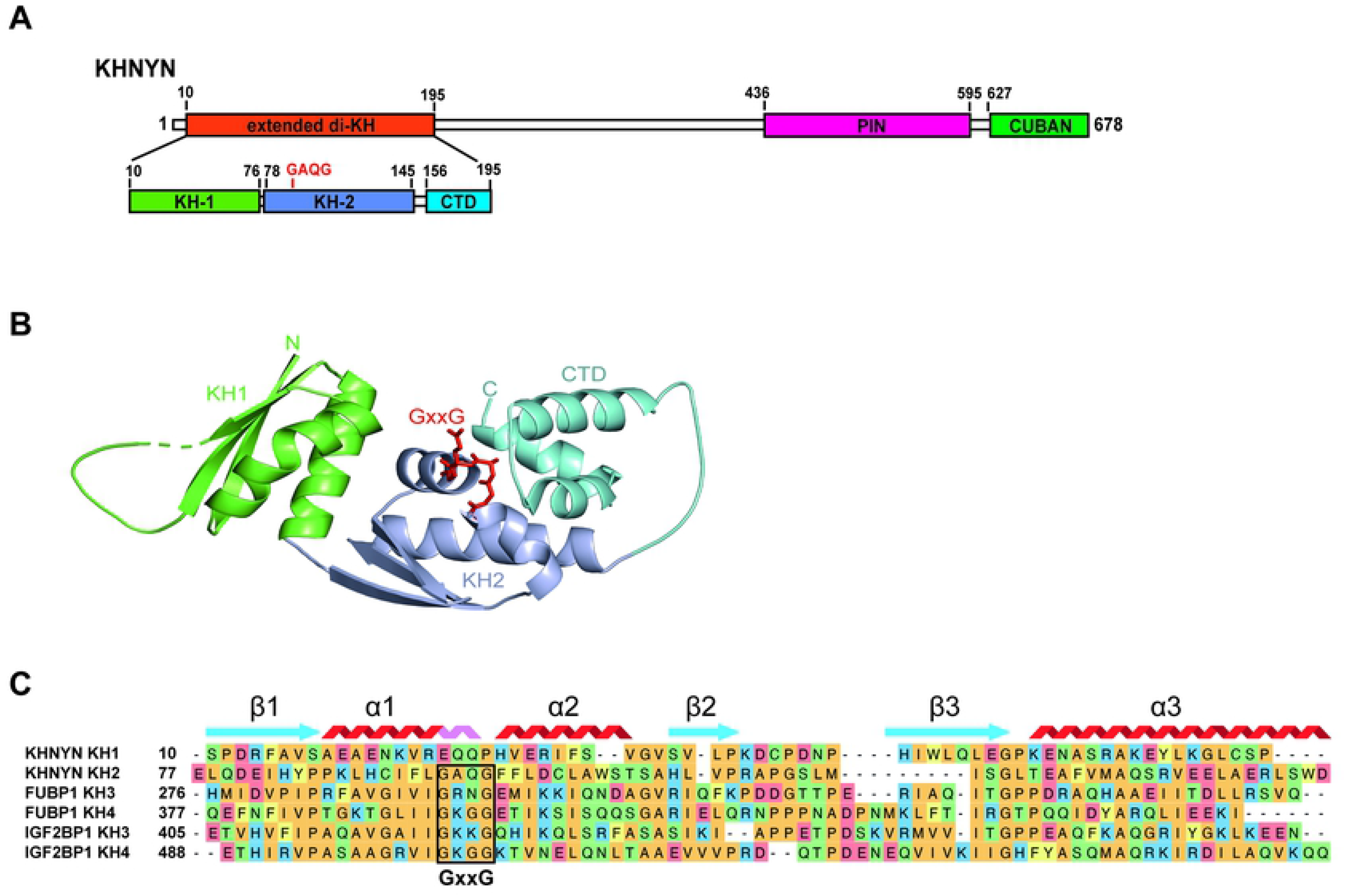
The extended di-KH domain in KHNYN. **(A)** A schematic diagram of KHNYN showing the extended di-KH domain, PIN domain and CUBAN domain. **(B)** Model for the extended di-KH domain. KH1, KH2 and C-terminal three-helix bundle are shown in cartoon representation colored green, blue and cyan, respectively. The GxxG motif in KH2 is highlighted in stick representation in red. **(C)** Sequence alignment of KH1 and KH2 from the KHNYN model, FUBP1 KH3 and KH4 (PDB ID: 1J4W) and IGF2BP1 KH3 and KH4 (PDB ID: 2N8L). The GxxG motif between α1 and α2 that is absent in KHNYN KH1 is boxed in the other sequences. The β1α1α2β2β3α3 secondary structure elements for the consensus KH domain based on 3D-alignment of the structures are displayed above the aligned sequences in light blue and red. The elongated portion of α1 present in KHNYN KH1 is shown in pink.

The top-ranked model from the Phyre2 analysis for KHNYN predicts a di-KH domain with a C-terminal extension from residues 10-195 based on the structure of the N4BP1 di-KH domain (Table S1). Of note, this differs with the previously published reports for KHNYN that described one KH domain [12, 13]. Models based on di-KH domains from the RNA binding proteins FUBP1, IGF2BP1 and KHSRP were also present in the analysis (Table S1). For the KHNYN di-KH domain model, KH1 is predicted from residues S10 to P76 while KH2 is predicted for residues L78 to D145 (Figure 1A-C). Type I KH domains consist of three β-strands that form a β-sheet and three α-helices that pack onto this surface [39]. Specifically, the sub-domain structure is β1-α1-α2-β2 followed by a variable loop and β3-α3 (Figure 1C). A hallmark GxxG loop between α1 and α2 forms one side of a hydrophobic RNA binding groove with these two α-helices while the β-sheet and the variable loop form the other side of the groove. Up to four nucleotides can be specifically recognized by a KH domain and the phosphates of the first two nucleotides interact with the GxxG loop. KH domains without the GxxG loop do not bind nucleic acid [39].

The KHNYN di-KH domain model shows two major differences with conventional di-KH domains. First, for KH1, the GxxG motif is not present between α1-α2 (Figure 1C). Instead, α1 appears to be elongated compared to that in FUBP1 or IGF2BP1 [40, 41]. Second, while the GxxG loop is present in KH2, an additional C-terminal three α-helix bundle (residues Q156 to Q195, cyan in Figure 1B) forms a sub-domain in which α2 and α3 of this module packs against α1, α2 and α3 of the KH2 module (blue in Figure 1B). Of note, the model for the elongated di-KH domain is similar to the AlphaFold model but does not support the previously proposed metal chelating module [13, 38]. In addition, the three-helix bundle extension has not been previously observed in other di-KH domain structures [40–44]. An uncharacterized domain previously named CGIN1 has been found only in KHNYN, N4BP1 and NYNRIN and this roughly correlates with the di-KH domain plus C-terminal extension [16]. Therefore, it appears that the KHNYN/N4BP1/NYNRIN family has a novel domain related to a di-KH domain but with potentially important unique characteristics and herein we refer to this as the extended di-KH domain.

In contrast to the extended di-KH domain identified by Phyre2, the SUPERFAMILY database annotation for KHNYN in Ensembl [45, 46] indicates only one type I KH-domain extending from residues 58-141. When we previously deleted this region, KHNYN had reduced antiviral activity [12]. However, this mutant protein localized in cytoplasmic puncta that were not visible for the wild-type protein, suggesting that this deletion may have altered the protein’s folding and led to its aggregation. To determine if deleting the extended di-KH domain affects KHNYN antiviral activity, we made a KHNYNΔdi-KH construct (Figure 2A) and analyzed its ability to restrict wild-type and CpG-enriched HIV-1. Of note, these experiments were performed in cells with endogenous KHNYN knocked out by CRISPR-Cas9-mediated genome engineering to eliminate the possibility that expression of the endogenous protein could affect the activity of the ectopically expressed protein, such as through multimerization. As previously shown [12], CRISPR-resistant wild-type KHNYN potently restricted HIV-1 with 36 CpGs introduced through synonymous mutations in the 5’ end of *env* (HIV-1_CpG_) (Figure 2B and Figure S1). However, deletion of the extended di-KH domain substantially reduced KHNYN antiviral activity for this virus. KHNYNΔdi-KH is expressed at moderately higher levels than wild-type KHNYN and is localized in the cytoplasm similar to wild-type KHNYN (Figure 2B and 2C), indicating that differences in its expression or subcellular localization do not account for its decreased antiviral activity.

**Figure 2.**
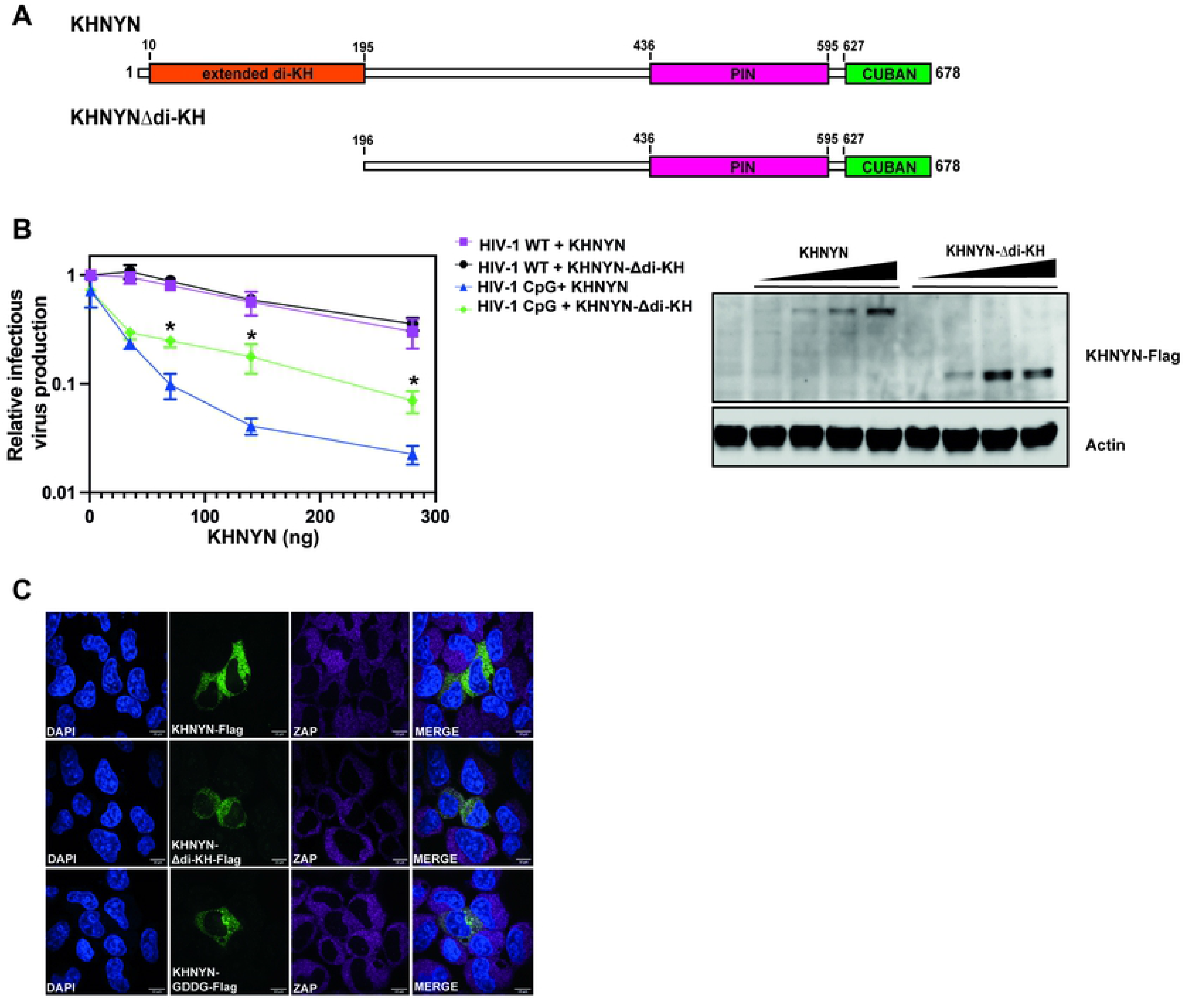
The extended di-KH domain is necessary for KHNYN antiviral activity against HIV-1_CpG_. **(A)** Schematic representation of KHNYNΔdi-KH. **(B)** Left panel: Infectious virus produced from KHNYN CRISPR HeLa cells co-transfected with HIV-1_WT_ or HIV-1_CpG_ and increasing concentration of CRISPR-resistant wild-type pKHNYN-FLAG or pKHNYNΔdi-KH-FLAG. Data are represented as mean ± standard deviation. Each point shows the average value of three independent experiments normalized to the value obtained for pHIV-1_WT_ at 0 ng pKHNYN. *p < 0.05 as determined by a two-tailed unpaired t-test comparing wild-type KHNYN and the mutant KHNYN at each concentration in the HIV-1_CpG_ samples. Right panel: representative western blot of wild-type KHNYN-FLAG or KHNYN-Δdi-KH-FLAG protein levels at concentrations corresponding to the left panel. **(C)** Representative confocal microscopy images of HeLa cells transfected as above and stained with an anti-FLAG antibody (green), anti-ZAP antibody (magenta) and DAPI (blue). The scale bar represents 10 μm.

### The extended di-KH domain in KHNYN does not mediate RNA binding

While canonical KH domains with a GxxG motif are found in many RNA binding proteins, a few proteins have a KH fold without a GxxG motif and do not bind RNA [39, 47, 48]. Since the GxxG loop in the extended di-KH domain is absent in KH1 but present in KH2, we tested whether KHNYN may bind RNA through KH2. One method for characterizing the RNA binding capacity of a KH domain is to make a rationally designed mutation that impairs RNA binding without altering the domain’s structure. This has typically been undertaken through the introduction of acidic residues between the glycine residues of the GxxG motif and has been shown to be a tool to probe whether a KH domain can bind RNA [49]. Therefore, we made a GAQG to GDDG mutation in KH2 (KHNYN-GDDG, Figure 3A). Interestingly, there was no loss of antiviral activity for KHNYN-GDDG compared to the wild-type protein for HIV-1CpG (Figure 3B and Figure S2). KHNYN-GDDG localized to the cytoplasm similar to wild-type KHNYN (Figure 2C). Therefore, while the extended di-KH domain is required for full antiviral activity, it may not bind RNA because KH1 does not contain a GxxG motif and mutating this motif in KH2 does not affect its ability to restrict HIV-1_CpG_.

**Figure 3.**
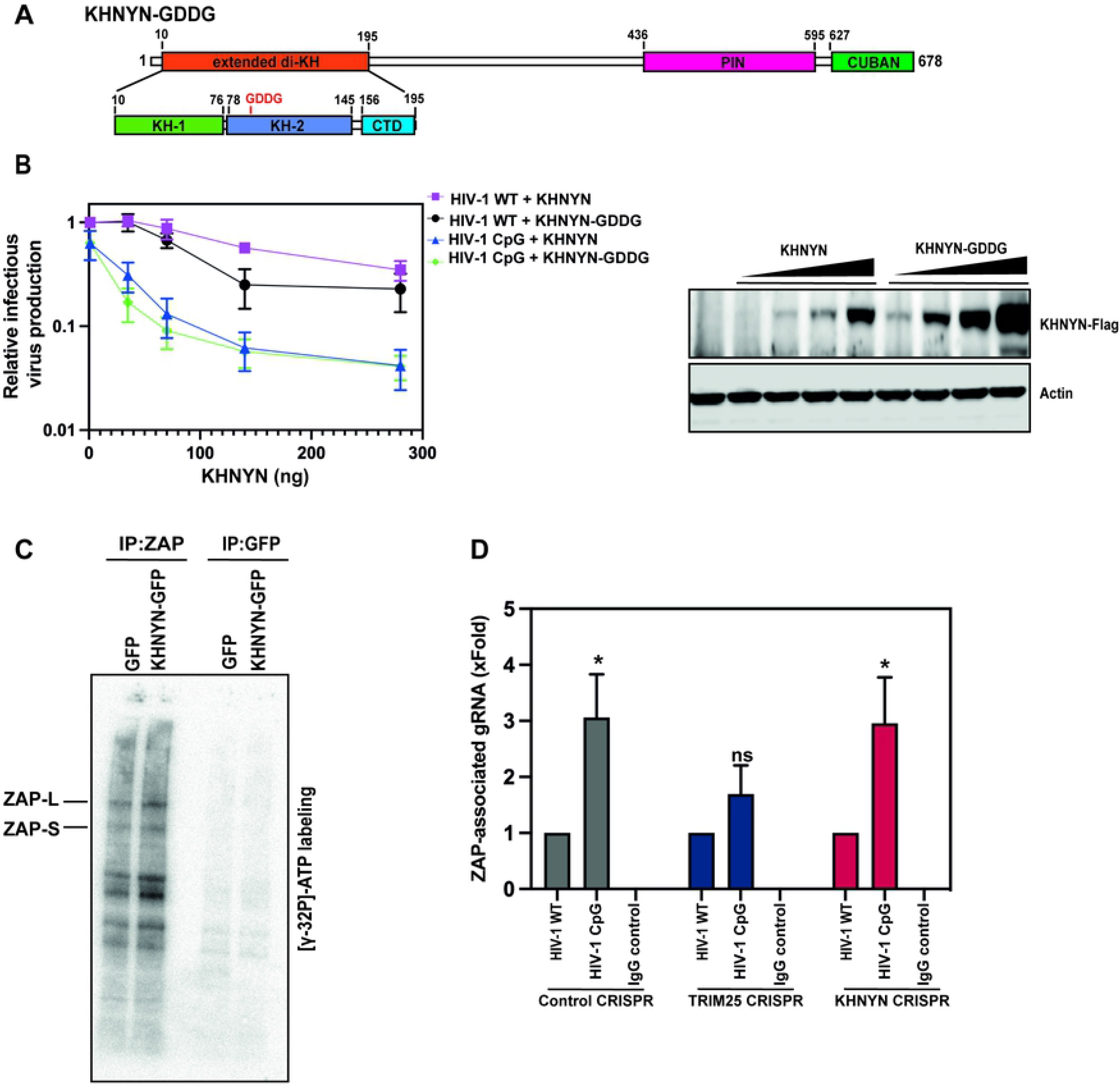
KHNYN does not detectably bind RNA and does not modulate ZAP binding to HIV-1 RNA. **(A)** Schematic representation of KHNYN-GDDG. **(B)** Left panel: Infectious virus produced from KHNYN CRISPR HeLa cells co-transfected with HIV-1_WT_ or HIV-1_CpG_ and increasing concentration of CRISPR-resistant wild-type pKHNYN-FLAG or pKHNYN-GDDG-FLAG. Data are represented as mean ± standard deviation. Each point shows the average value of three independent experiments normalized to the value obtained for pHIV-1_WT_ at 0 ng pKHNYN. Right panel: representative western blot of wild-type KHNYN-FLAG or KHNYN-GDDG-FLAG protein levels at concentrations corresponding to the left panel. **(C)** SDS-PAGE gel of γ^32^P-RNA labelled proteins from KHNYN CRISPR HeLa cells stably expressing GFP or wild-type KHNYN-GFP. After UV crosslinking and cell lysis, immunoprecipitations were performed using antibodies targeting GFP or endogenous ZAP. The background bands are RNA binding proteins that co-immunoprecipitate with ZAP. **(D)** Quantification of HIV-1 genomic RNA (gRNA) bound to ZAP after UV crosslinking and immunoprecipitation in control CRISPR cells, TRIM25 CRISPR cells or KHNYN CRISPR cells. Each bar shows the average value of three independent experiments normalized to the value obtained for HIV-1_WT_ in each cell line. *p < 0.05 as determined by a two-tailed unpaired t-test comparing HIV-1_CpG_ to HIV-1_WT_ in each cell line.

KHNYN is not listed as a human RNA binding protein in RBPbase [50], though this could be due to its low expression level [51]. Quantitative proteomics have shown that many of the characterized ZAP cofactors are expressed at much higher levels than KHNYN (Figure S3A) and there are ~25-fold more ZAP molecules/cell than KHNYN in HeLa cells [51]. To determine if KHNYN directly bound RNA, we analyzed whether RNA could be crosslinked by ultraviolet (UV) light to KHNYN in stable cell lines that express CRISPR-resistant KHNYN-GFP in KHNYN knockout cells. These cells restricted HIV-1_CpG_ and expressed KHNYN at a higher level than the endogenous protein (Figure S3B-C). Briefly, cells expressing GFP or KHNYN-GFP were UV-C (254 nm) crosslinked, lysed and either ZAP or GFP were immunoprecipitated. The samples were treated with RNase I, which cleaves single-stranded RNA non-specifically and so degrades protein-bound RNA bar a few nucleotides. The remaining crosslinked RNA was labelled with γ^32^P-ATP, the complexes resolved by SDS-PAGE and visualized using a Phosphorimager after transfer to a nitrocellulose membrane. The autoradiograph shows two bands in the ZAP immunoprecipitates corresponding to the size of ZAP-L and ZAP-S, demonstrating that, as expected, both proteins bind RNA directly (Figure 3C). Additional bands on the autoradiograph for the ZAP immunoprecipitates presumably correspond to RNA binding proteins that co-purify with ZAP. In contrast, even though KHNYN is well expressed (Figure S3C), it does not show specific radiolabeling in the same samples, indicating that it does not stably bind RNA (Figure 3C).

To determine if KHNYN is required for ZAP to bind to CpG-enriched HIV-1, we immunoprecipitated endogenous ZAP in control and KHNYN knockout HeLa cells and determined the abundance of the co-purifying HIV-1 RNA using quantitative RT-PCR. As a control, we also tested the role of the ZAP cofactor TRIM25, which has previously been shown to be required for optimal ZAP binding to a target RNA with a ZAP-response element from murine leukemia virus [10]. ZAP binds HIV-1_WT_ RNA but the amount of viral RNA associated with ZAP is higher for HIV-1_CpG_ (Figure 3D). This is consistent with the increased binding of ZAP to the clustered CpGs in HIV-1_CpG_ *env* observed in UV crosslinking and immunoprecipitation sequencing (CLIP-seq) experiments [28]. As expected, ZAP enrichment on HIV-1_CpG_ compared to HIV-1_WT_ is reduced in TRIM25 knockout cells. However, there were similar levels of enrichment for ZAP binding to HIV-1_CpG_ RNA compared to HIV-1_WT_ RNA in the control and KHNYN knockout cells, implying that KHNYN does not affect ZAP binding to CpG-enriched HIV-1 RNA. Because ZAP is a type I and II interferon-stimulated gene [52, 53], we also tested whether KHNYN was required for ZAP binding to HIV-1_CpG_ when ZAP-L was overexpressed. Similar to the result observed at endogenous ZAP levels, KHNYN depletion had no effect for ZAP enrichment on HIV-1_CpG_ RNA (Figure S3D). This is consistent with our previous results showing that the sensitivity of different CpG-enriched HIV-1 genomes to ZAP determines the degree by which they are restricted by KHNYN [29]. Overall, this suggests that KHNYN is not required for ZAP to bind HIV-1_CpG_ and acts at a later point in the restriction pathway.

### The CUBAN domain in KHNYN regulates its abundance and subcellular localization

The C-terminal CUBAN domain in KHNYN (Figure 4A) has been shown to bind both NEDD8 and ubiquitin, with preferential binding to monomeric NEDD8 over ubiquitin [15]. This mediates an interaction between KHNYN and the neddylated cullin-RING E3 ubiquitin ligases, including CUL1, CUL2, CUL3 and CUL4, and raised the possibility that KHNYN antiviral activity could be regulated by these proteins. To determine whether the CUBAN domain is required for KHNYN antiviral activity, it was deleted (KHNYNΔCUBAN, Figure 4A) and the ability to restrict HIV-1CpG was measured. Deletion of the CUBAN domain led to a large decrease in KHNYN antiviral activity, even though KHNYNΔCUBAN was expressed at much higher levels than the wild-type protein (Figure 4B and Figure S4A). This indicates that the domain may be required for both KHNYN homeostatic turnover and antiviral activity.

**Figure 4.**
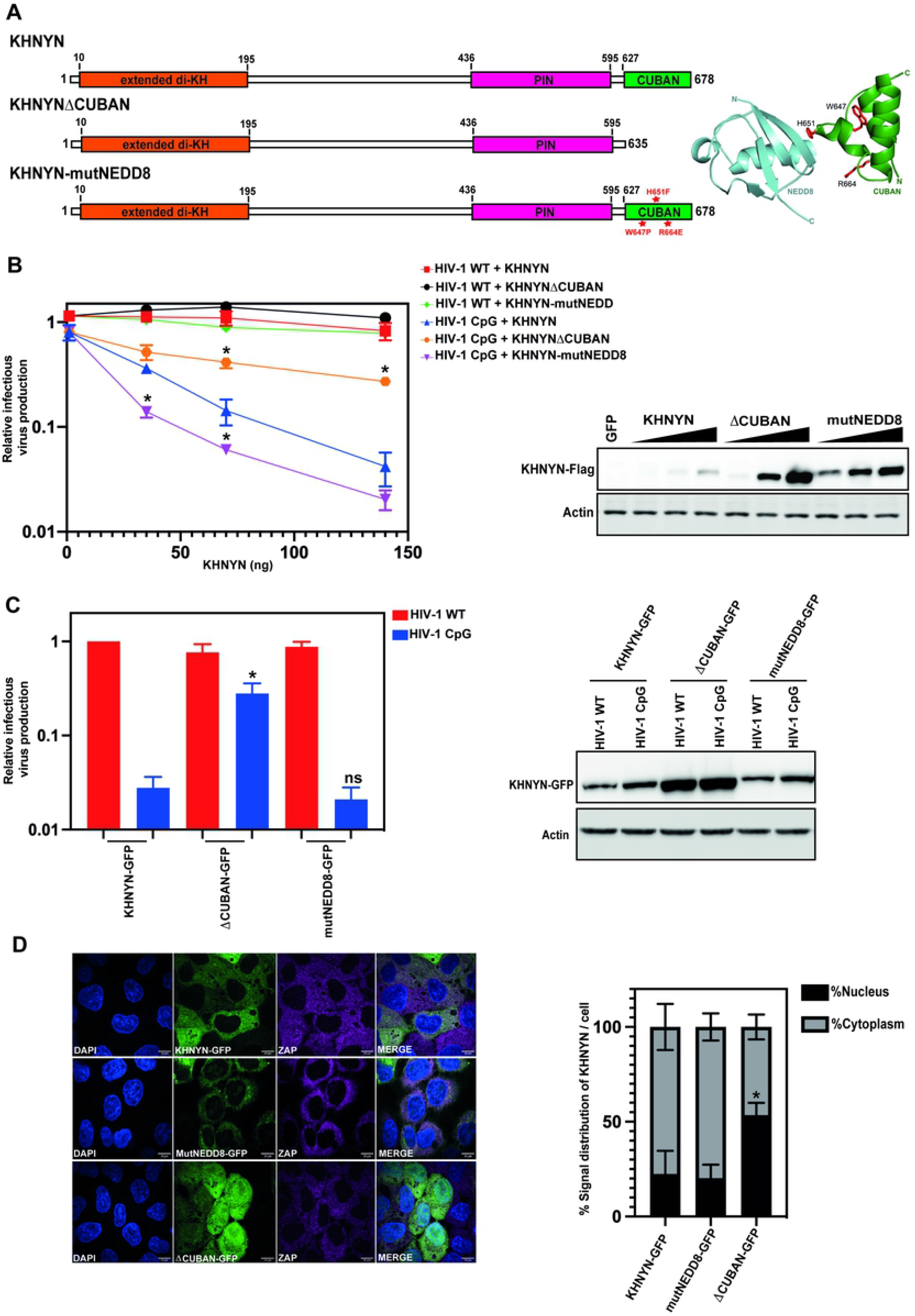
The CUBAN domain is essential for KHNHN antiviral activity and protein localization. **(A)** Left panel: Schematic representation of KHNYNΔ636-678 (KHNYNΔCUBAN) and KHNYN-W647P/H651F/R664E (KHNYNmutNEDD8). The residues mutated in KHNYNmutNEDD8 are highlighted in red. Right panel: Cartoon representation of the NEDD8-CUBAN complex structure. The residues mutated in KHNYNmutNEDD8 are highlighted in stick representation in blue. **(B)** Left panel: Infectious virus production from KHNYN CRISPR HeLa cells co-transfected with pHIV-1_WT_ or pHIV-1_CpG_ and increasing concentration of CRISPR-resistant wild-type pKHNYN-Flag, pKHNYNΔCUBAN-Flag or pKHNYN-mutNEDD8-Flag. Each point shows the average value of three independent experiments normalized to the value obtained for pHIV-1_WT_ at 0ng pKHNYN. *p < 0.05 as determined by a two-tailed unpaired t-test comparing wild-type KHNYN and the mutant KHNYN at each concentration in the HIV-1_CpG_ samples. Right panel: representative western blot of the protein level of wild type KHNYN-FLAG, KHNYNΔCUBAN-FLAG and KHNYNmutNEDD8-FLAG at the concentrations shown in the left panel. **(C)** Left panel: Infectious virus production from Hela KHNYN CRISPR cells stably expressing wild-type KHNYN-GFP, KHNYNΔCUBAN-GFP or KHNYNmutNEDD8-GFP. All cell lines were transfected with HIV-1_WT_ or HIV-1_CpG_. Each bar shows the average value of five independent experiments normalized to the value obtained for wild type KHNYN co-transfected with pHIV-1_WT_. *p < 0.05 as determined by a two-tailed unpaired t-test comparing wild-type KHNYN and the mutant KHNYN construct in the HIV-1_CpG_ samples. Data are represented as mean ± SD. Right panel: Representative western blot for GFP showing the KHNYN-GFP protein levels in the wild-type KHNYN-GFP, KHNYNΔCUBAN-GFP or KHNYNmutNEDD8-GFP cell lines. **(D)** Left panel: Confocal microscopy images of the KHNYN-GFP cell lines (green) co-stained with endogenous ZAP (magenta), scale bar is 10μm. Right panel: Signal quantification per cell (50 cells total per condition) of KHNYN nuclear and cytoplasmic distribution in the KHNYN-GFP cell lines. *p < 0.05 as determined by a two-tailed unpaired t-test comparing the nuclear fraction between each sample.

The CUBAN domain comprises a three-helix bundle (α1 = T632-R640, α2 = K652-L657, α3 = Y662-F678) connected by two loops [15]. The binding interface between the CUBAN domain and NEDD8 is formed from a negatively charged motif in NEDD8 and a positively charged surface in the CUBAN α2 and surrounding residues. Three mutations (Figure 4A) have previously been shown to decrease binding of the CUBAN domain to NEDD8 [15]. H651F and R664E decrease the electrostatic binding interaction while W647P likely increases the flexibility of loop 1 (Q641-H651). Surprisingly, when these mutations were introduced in KHNYN (KHNYN-mutNEDD8), they moderately increased its antiviral activity for HIV-1_CpG_ (Figure 4B and Figure S4A). This correlated with increased protein abundance (Figure 4B), suggesting that the NEDD8 interaction likely regulates KHNYN turnover but not antiviral activity.

To analyze the subcellular localization of the mutant KHNYN proteins, KHNYN-mutNEDD8-GFP and KHNYNΔCUBAN-GFP stable cell lines were made. Similar to the transient transfection experiments described above (Figure 4B), deletion of the CUBAN domain decreased KHNYN-GFP antiviral activity while introducing the mutations that reduce NEDD8 binding did not affect it (Figure 4C and Figure S4B). Of note, the increase in KHNYN abundance due to the CUBAN domain deletion or mutations that decrease NEDD8 binding are less pronounced in the KHNYN-GFP cell lines than in the experiments with transiently transfected KHNYN-FLAG constructs, possibly because the GFP fusion stabilizes the wild-type protein. Interestingly, while KHNYN-mutNEDD8-GFP had a similar localization to the cytoplasm as wild-type KHNYN-GFP, KHNYNΔCUBAN-GFP had a substantial increase in nuclear localization (Figure 4D). Therefore, the CUBAN domain regulates KHNYN subcellular localization in addition to its homeostatic turnover and antiviral activity.

### KHNYN has a nuclear export signal at the C-terminus of the CUBAN domain that is required for antiviral activity

CRM1 (also known as XPO1) is a nuclear export protein that mediates trafficking of a large number of cellular proteins and ribonucleoprotein complexes from the nucleus to the cytoplasm [54]. CRM1 binds leucine-rich nuclear export signals (NESs) in cargo proteins and KHNYN has previously been identified as a CRM1 cargo in a large-scale proteomics screen [54, 55]. To confirm that KHNYN uses the CRM1 nuclear export pathway, we compared the subcellular localization of KHNYN-GFP, KHNYN-mutNEDD8-GFP and KHNYNΔCUBAN-GFP in the absence and presence of leptomycin B, a small molecule inhibitor of CRM1 [56]. Addition of leptomycin B to the KHNYN-GFP stable cell lines substantially increased wild-type KHNYN-GFP or KHNYN-mutNEDD8-GFP nuclear localization to levels similar to KHNYNΔCUBAN-GFP (Figure 5). However, the subcellular localization of KHNYNΔCUBAN-GFP was not affected by leptomycin B, supporting the observation that the functional NES for CRM1 is present in the CUBAN domain. ZAP-L and ZAP-S also have a CRM1 NES and ZAP-S has been reported to be a CRM1-dependent nucleocytoplasmic shuttling protein [55, 57]. However, ZAP was not re-localized to the nucleus by leptomycin B treatment (Figure 5), which may indicate that in these cells it is sequestered in the cytosol or at cytoplasmic membranes.

**Figure 5.**
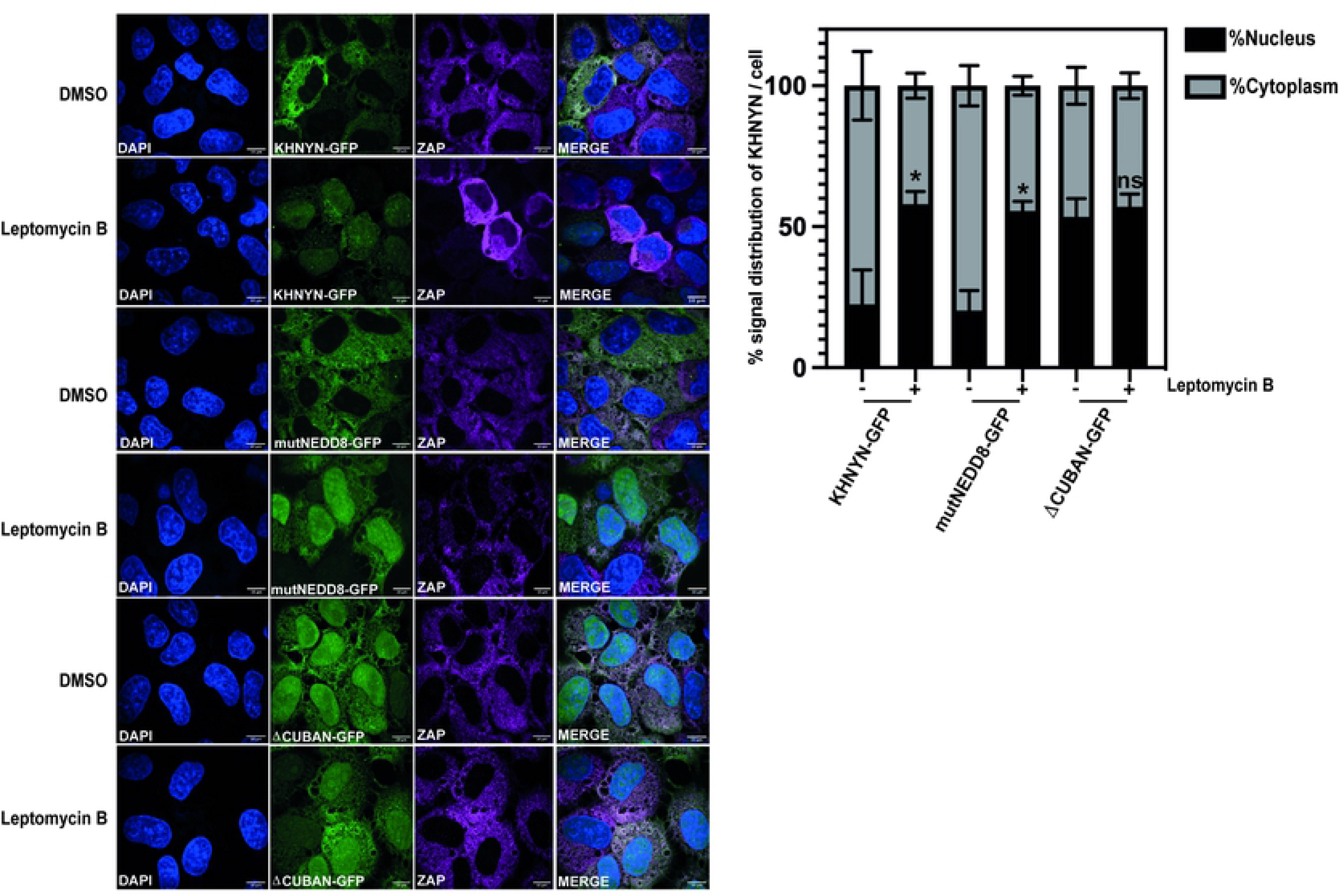
CRM1 inhibition re-localizes KHNYN to the nucleus. Left panel: Representative confocal microscopy images of KHNYN CRISPR HeLa cells stably expressing either wild-type KHNYN-GFP or the indicated mutants before and after a four-hour treatment with 50nM Leptomycin B. KHNYN-GFP is shown in green, endogenous ZAP co-staining is shown in magenta, scale bar is 10μm. Right panel: Signal quantification per cell (50 cells in total per condition) of KHNYN nuclear and cytoplasmic distribution for KHNYN-GFP or the indicated mutant protein before and after Leptomycin B treatment. *p < 0.05 as determined by a two-tailed unpaired t-test comparing the nuclear fraction between each sample.

There are at least two types of CRM1 NESs with different spacing of the hydrophobic residues that fit into five pockets in CRM1: the Rev-type NES and the PKI-type NES [58]. To identify potential NESs in KHNYN, we used the Wregex tool [59], which identified a putative NES at the C-terminus of the CUBAN domain with hydrophobic residues fitting the PKI-type NES spacing (residues 669-678, <u>L</u>SEA<u>L</u>LS<u>L</u>N<u>F</u>, amino acids predicted to bind CRM1 are underlined). Of note, this tool only identified positions 1-4 for the predicted CRM1 NES and did not identify the more recently identified position 0 [58, 59]. For a PKI-type NES, position 0 is two amino acids upstream of position 1 and is preferentially preceded by an acidic residue [58]. The C-terminal NES in KHNYN fits this consensus perfectly, with the full NES predicted to be D<u>I</u>NQ<u>L</u>SEA<u>L</u>LS<u>L</u>N<u>F</u> and comprising the last 14 amino acids of the protein (Figure 6A). This sequence is located in the third helix of the CUBAN domain (Figure 6B), which does not contain any of the residues that interact with NEDD8.

**Figure 6.**
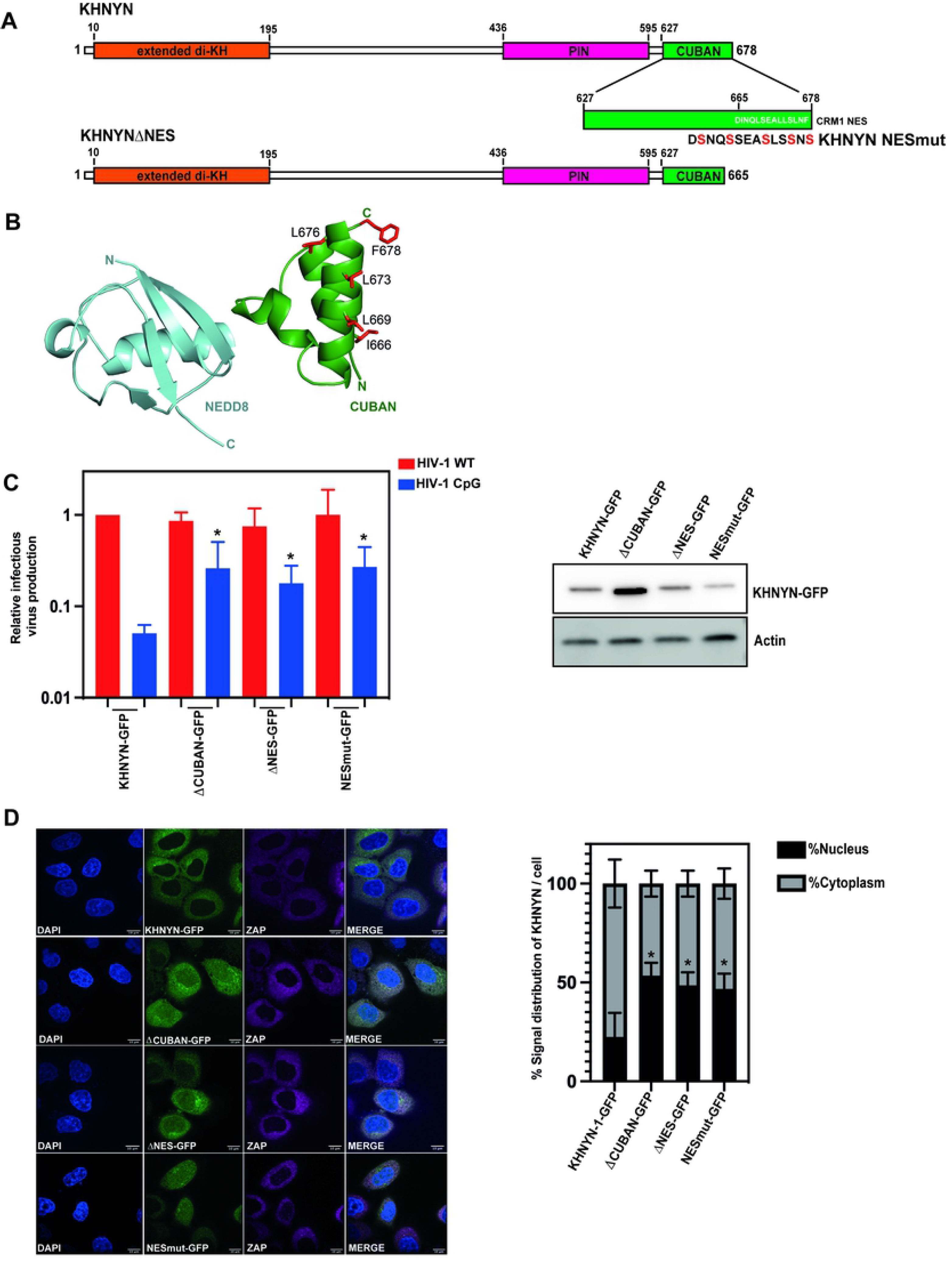
A nuclear export signal present in the C-terminal 14 amino acids of the CUBAN domain is required for KHNYN cytoplasmic location and antiviral activity. **(A)** Schematic representation of the KHNYN CUBAN domain and nuclear export signal. The key residues that were mutated in KHNYN-NESmut are shown in red. **(B)** Cartoon representation of the NEDD8-CUBAN complex structure. Residues in the NES that were mutated to serine are shown as sticks in red. The mutations in KHNYNmutNEDD8 that decrease KHNYN binding to NEDD8 are shown as sticks in blue. **(C)** Left panel: Infectious virus production from KHNYN CRISPR HeLa cells stably expressing wild-type KHNYN-GFP, KHNYNΔCUBAN-GFP, KHNYNΔNES-GFP or KHNYN-NESmut-GFP transfected with HIV-1_WT_ or HIV-1_CpG_. Each bar shows the average value of three independent experiments normalized to the value obtained for wild type KHNYN co-transfected with pHIV-1_WT_. *p < 0.05 as determined by a two-tailed unpaired t-test comparing each mutant KHNYN-GFP sample to wild type KHNYN-GFP. Right panel: Representative western blot showing KHNYN-GFP protein levels. **(D)** Left panel: Confocal microscopy images of the KHNYN-GFP cell lines (green) co-stained for endogenous ZAP (magenta), scale bar is 10μm. Right panel: Signal quantification per cell (50 cells total per condition) of KHNYN nuclear and cytoplasmic distribution in the KHNYN-GFP cell lines. *p < 0.05 as determined by a two-tailed unpaired t-test comparing the nuclear fraction between each sample.

To test the functional role of the NES in KHNYN, we made stable cell lines expressing KHNYNΔNES-GFP and KHNYN-NESmut-GFP. KHNYN-NESmut-GFP has all five amino acids predicted to directly bind CRM1 mutated to serine and, in KHNYNΔNES-GFP, the entire 14 residue NES sequence was deleted (Figure 6A). KHNYNΔNES-GFP does not show a similar increase in protein abundance as KHNYNΔCUBAN-GFP and has similar expression levels as KHNYN-GFP (Figure 6C), indicating that the homeostatic turnover mediated by the NEDD8 interaction is not affected by deleting the final 14 amino acids of the domain. It should be noted that the three-helix bundle of the CUBAN domain fold is not tightly packed and NEDD8 binding may not require the integrity of the fully folded domain [15]. Deleting or mutating the NES decreased KHNYN antiviral activity and increased its nuclear localization similar to KHNYN-ΔCUBAN (Figure 6C-D and Figure S5A). This suggests that the loss of antiviral activity for KHNYNΔCUBAN is due to the deletion of the C-terminal NES in the CUBAN domain. This also shows that changes in KHNYN abundance can be separated from changes in its subcellular localization and antiviral activity. While deletion or mutation of the NES re-localizes KHNYN to the nucleus, it has no effect on ZAP or TRIM25 localization (Figure 6D and Figure S5B).

Because we did not observe ZAP re-localization to the nucleus upon CRM1 inhibition by leptomycin B treatment or re-localization of KHNYN to the nucleus (Figure 4D, Figure 5 and Figure 6D), we hypothesized that the KHNYN NES could be required for antiviral activity by allowing it to interact with ZAP in the cytoplasm. ZAP-L has greater antiviral activity than ZAP-S against several viruses, including HIV-1_CpG_ [32–35]. This is at least in part because it contains an S-farnesylation motif that localizes it to the endomembrane system and is required for its antiviral activity [33–36]. To determine if the nuclear export signal in KHNYN is required for it to interact with ZAP, we performed co-immunoprecipitation experiments in the KHNYN-GFP and KHNYNΔNES-GFP cell lines. Immunoprecipitation of KHNYN-GFP pulled down

ZAP-L but no detectable ZAP-S (Figure 7A). Supporting this observation, we recently showed that immunoprecipitation of ZAP-L preferentially pulls down KHNYN compared to ZAP-S and mutation of the S-farnesylation motif decreases this interaction [35]. KHNYNΔNES-GFP immunoprecipitated less ZAP-L than wild-type KHNYN (Figure 7A), which suggests that nuclear export of KHNYN is required for it to bind ZAP-L that is localized to the cytoplasmic endomembrane system.

**Figure 7.**
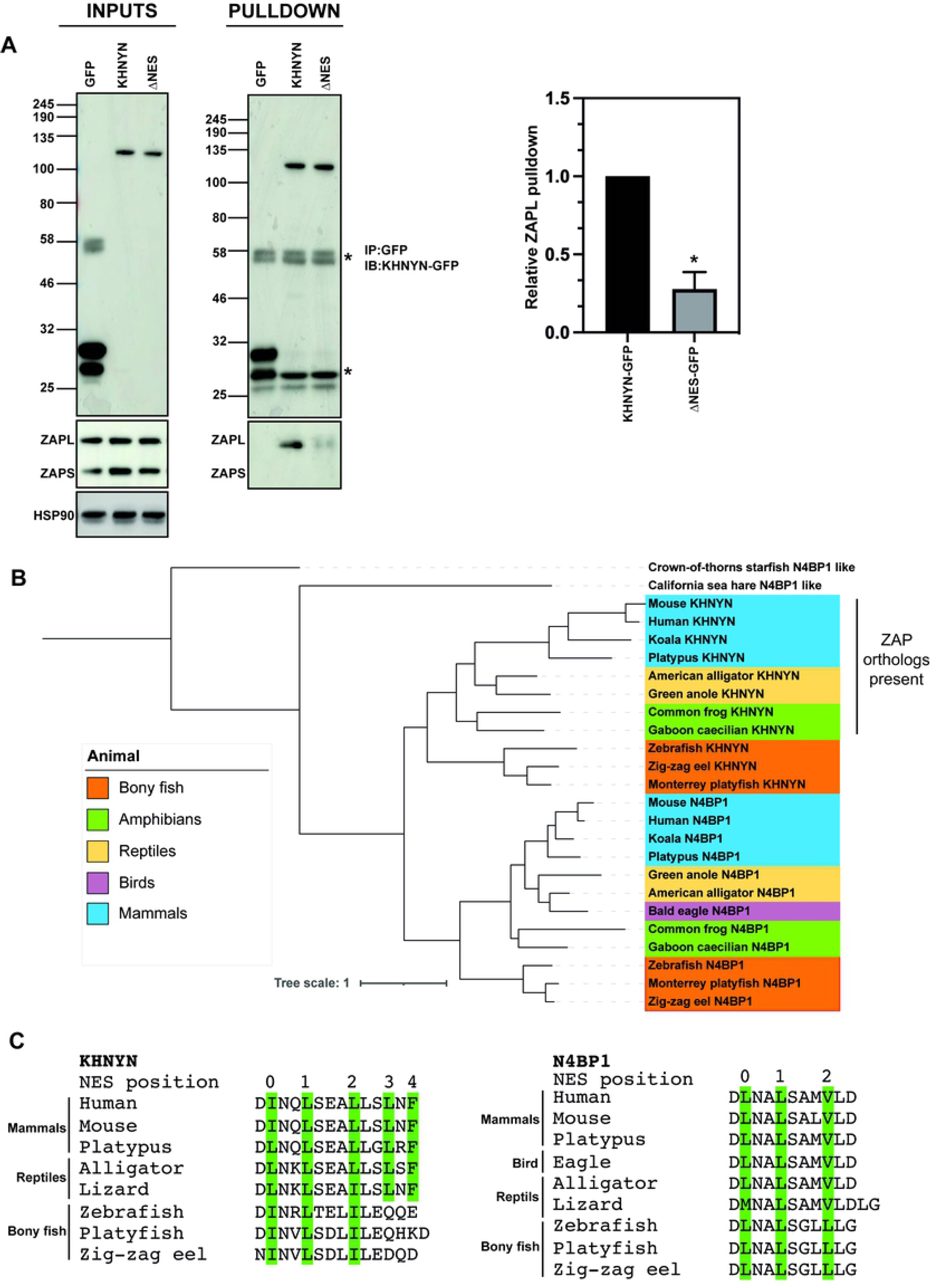
The KHNYN NES in the CUBAN domain promotes its interaction with ZAP and evolved at a similar time as ZAP in tetrapods. **(A)** Left panel: GFP control, KHNYN-GFP and KHNYNΔNES-GFP HeLa cells were lysed and aliquots were immunoblotted for GFP and endogenous ZAP. Middle panel: GFP was immunoprecipitated in each cell lysate and blotted for GFP or endogenous ZAP. Right panel: The amount of ZAP-L immunoprecipitated relative to the KHNYN-GFP sample is presented in a bar graph, N = 3. * p = <0.05 as determined by a two-tailed unpaired t-test. **(B)** Maximum likelihood phylogenetic tree of KHNYN and N4BP1 amino acid sequences. Representative sequences from bony fish (orange), amphibians (green), reptiles (yellow), birds (light purple) and mammals (light blue) were aligned and maximum likelihood phylogeny was inferred using the LG substitution model DIVEIN. The crown-of-thorns starfish and Californian sea hare were used as outgroups to root the tree. An unpruned phylogenetic tree including all analysed sequences, their scientific names and sequence accession numbers is presented in Figure S6. **(C)** Alignment of the C-terminal NES residues of KHNYN and N4BP1 from selected species. Residues in PKI-like NES positions 0 – 4 are highlighted in green.

ZAP and TRIM25 have previously been shown to have co-evolved [60] and we hypothesized that KHNYN might also have acquired adaptive changes to act as a ZAP cofactor. ZAP evolved in tetrapods from a PARP12 gene duplication [60]. While the N4BP1-KHNYN evolutionary pathway is currently unclear, N4BP1-like proteins are present outside of chordates such as in the crown-of-thorns starfish (*Acanthaster planci*). Within the phylum *Chordata*, N4BP1 proteins are found throughout the *Vertebrata* subphylum, including bony fish (class *Osteichthyes*) (Figure 7B and Figure S6). Clear KHNYN orthologs are also present in bony fish and are present in most tetrapods, including amphibians (class *Amphibia*), reptiles (class *Reptilia*) and mammals (class *Mammalia*). However, while N4BP1 orthologs are present in birds (class *Aves*), KHNYN orthologs are not, suggesting that it has been lost in this lineage. Interestingly, the evolution of the KHNYN NES appears to correlate with the evolution of ZAP in tetrapods. In bony fish, the C-terminus of KHNYN contains three hydrophobic amino acids in positions 0, 1 and 2 of the NES (Figure 7C). These are also present in N4BP1 orthologs throughout vertebrates. However, for the amphibian, reptile and mammal classes with clear ZAP orthologs [60], hydrophobic residues are found in positions 3 and 4 of the NES and are conserved throughout these lineages (Figure 7C and Figure S6). Therefore, we hypothesize that the NES at the C-terminus of KHNYN co-evolved with the evolution of ZAP from PARP12 in the tetrapod lineage.

## DISCUSSION

KHNYN has three primary domains: an N-terminal extended di-KH domain, a PIN domain and a C-terminal CUBAN domain. All three domains are necessary for its antiviral activity. The extended di-KH domain appears to be unique to the KHNYN/N4BP1/NYNRIN family and has diverged from previously characterized di-KH domains in RNA binding proteins. In KHNYN, the GxxG motif required for RNA binding is not present in KH1. Furthermore, the di-KH domain is extended by the packing of three additional alpha helices at the C-terminus of KH2, forming what may be a functional module. While the extended di-KH domain is required for full antiviral activity, our data shows that it does not appear to stably bind RNA since there was no detectable crosslinking of KHNYN to cellular RNA and it does not modulate ZAP binding to CpG-enriched HIV-1 RNA. KH domains in the Rrp40 or GLD-3 proteins do not bind RNA and instead have been shown to mediate protein-protein interactions [47, 48]. This is a potential function for the extended di-KH domain in KHNYN. Of note, the ZAP interaction region in KHNYN identified through a yeast-two-hybrid screen consists of the C-terminal portion of the PIN domain plus the CUBAN domain, so the extended di-KH domain likely does not mediate the ZAP-KHNYN interaction [12]. However, it could regulate the binding of other, currently unidentified, ZAP cofactors.

The low abundance of KHNYN appears to be at least partly due to its CUBAN domain, which mediates an interaction with neddylated cullin-RING E3 ubiquitin ligases [15]. Mutation of the residues in this domain that bind NEDD8 lead to a substantial increase in KHNYN abundance, though this does not affect its antiviral activity. KHNYN protein levels could be tightly regulated to prevent off-target endoribonuclease activity, which would be determinantal to cellular gene expression, thus necessitating turnover by cullin-RING E3 ubiquitin ligases. In HeLa cells, ZAP is expressed at much higher levels than KHNYN [51]. This appears to make it limiting for ZAP antiviral activity because KHNYN overexpression potently promotes restriction of CpG-enriched HIV-1 [12]. In addition, KHNYN could have additional cellular functions beyond being a ZAP cofactor since KHNYN orthologs are present in bony fish that do not have clear ZAP orthologs.

It remains unclear how the ZAP antiviral complex is assembled on a ZAP-response element in viral RNA to inhibit gene expression [1]. While CLIP-seq experiments have shown that ZAP binds at least transiently to several CpG sites in wild-type HIV-1, this does not correlate with substantial antiviral activity [3, 28, 60]. Instead, a substantial number of CpG dinucleotides appear to have to be clustered together, possibly with specific context preferences such as CpG spacing, flanking nucleotide composition and local RNA structure, to allow ZAP to target HIV-1 RNA for degradation and restrict viral replication [28–30]. Several lines of evidence support that ZAP mediates KHNYN targeting to viral RNA. First, KHNYN is not required for ZAP to bind CpG-enriched HIV-1 RNA. Second, when ZAP is depleted, KHNYN loses its antiviral activity [12]. Third, HIV-1 with CpGs introduced in different regions of the genome have differential sensitivity to ZAP and this correlates with KHNYN antiviral activity [29]. Therefore, we propose that ZAP and KHNYN interact to form a complex in which ZAP provides the RNA targeting module and KHNYN cleaves the target RNA through its PIN domain, which we have previously shown to be required for antiviral activity [12]. TRIM25 appears to regulate ZAP binding to target HIV-1 RNA and therefore the presence of TRIM25 binding sites may also be important to define ZAP-response elements.

ZAP subcellular localization appears to regulate its antiviral activity against several viruses in that ZAP-L with an intact S-farnesylation motif mediates more potent restriction than ZAP-S [32–35]. This correlates with preferential KHNYN and TRIM25 binding to ZAP-L compared to ZAP-S, even though the binding sites for KHNYN and TRIM25 are present in both isoforms [9, 12, 35, 60]. KHNYN subcellular localization also appears to be important in that the CRM1 nuclear export signal is required for full antiviral activity. How KHNYN is targeted to the nucleus is not clear and we have not identified a canonical nuclear localization signal (NLS) in it. However, KHNYN could be trafficked into the nucleus by interacting with other proteins that contain an NLS. Where KHNYN first interacts with ZAP is also not known and ZAP-L localization may be dynamic. One possibility is that cytoplasmic ZAP-L molecules bind KHNYN before or after localization to the endomembrane system but prior to binding RNA, leading to a pre-formed antiviral complex. However, the limiting abundance of KHNYN implies that only a small pool of ZAP molecules would be bound to KHNYN. Another possibility is that KHNYN cycles through the nucleus and cytoplasm and only interacts with ZAP after it binds RNA, which would act as a regulatory mechanism to allow endonucleolytic cleavage only for RNAs that have ZAP bound to them with a particular stoichiometry or structure. Therefore, in addition to its low abundance, nuclear localization of KHNYN could regulate its activity by preventing it from interacting with ZAP bound RNAs that are not bona fide targets. Future experiments to define the molecular characteristics and subcellular localization of the ZAP-KHNYN antiviral complex in a living cell will be required to fully understand how ZAP inhibits viral gene expression.

## MATERIALS AND METHODS

### Plasmids and Cell lines

HeLa, HEK293T and TZM-bl cells were maintained in high glucose DMEM supplemented with GlutaMAX (Thermo Fisher Scientific), 10% fetal bovine serum, 100 U/mL penicillin and 100 mg/mL streptomycin and incubated with 5% CO2 at 37°C. Control CRISPR, KHNYN CRISPR and TRIM25 CRISPR HeLa cells were previously described [12]. The CRISPR-resistant pKHNYN-FLAG plasmid has been previously described [12] and specific mutations were cloned into it using site-specific mutagenesis. All primers (Supplementary Table 2) were purchased from Eurofins Genomics and all PCR reactions were performed using Q5 High-Fidelity (New England Biolabs). HIV-1_NL4-3_ (pHIV-1_WT_) and HIV*env*_86-561CpG_ (pHIV-1_CpG_) in pGL4 were previously described [12, 61]. CRISPR-resistant KHNYN-GFP constructs were made using a flexible "GGGGSGGGGSGGGG" linker between KHNYN and GFP. Stable CRISPR KHNYN HeLa cells expressing CRISPR-resistant KHNYN-GFP, KHNYNmutNEDD8-GFP, KHNYNΔCUBAN-GFP, KHNYNΔNES-GFP and KHNYN-NESmut-GFP were produced by transduction with the murine leukemia virus (MLV) retroviral vector MIGR1 with the KHNYN-GFP construct cloned into the multiple cloning site and GFP replaced by the Blasticidin S-resistance gene [62].

### Transfections and infections

HeLa cells were seeded in 24-well plates at 70% confluency. Cells were transfected according to the manufacturer’s instructions using TransIT-LT1 (Mirus) at the ratio of 3 μL TransIT-LT1 to 1 μg DNA. For the HIV experiments, 0.5 μg pHIV_WT_ or pHIV_CpG_ and the designated amount of KHNYN-FLAG or GFP-FLAG for a total of 1 μg DNA were transfected. 24 hours post-transfection, the culture media was replaced with fresh media. For HIV-1WT or HIV-1_CpG_ infection of HeLa cells, viral stocks were produced by co-transfecting pHIV-1_WT_ or pHIV-1_CpG_ with pVSV-G [63] into HEK293T ZAP CRISPR cells [29] and titred on TZM-bl cells.

### Analysis of protein expression by immunoblotting

48 hours post-transfection, the HeLa cells were lysed in Laemmli buffer at 95°C for 10 minutes. The culture supernatant was filtered through a 0.45 μM filter and virions were pelleted by centrifugation for 2 hours at 20,000 x g through a 20% sucrose cushion in phosphate-buffered saline (PBS). Viral pellets were resuspended in 2X Laemmli buffer. Cell lysates and virion lysates were resolved on 8 to 16% Mini-Protean TGX precast gels (Bio-Rad), transferred onto nitrocellulose membranes (GE Healthcare) and blocked in 5% non-fat milk in PBS with 0.1% Tween 20. Primary antibodies were incubated overnight at 4°C followed by 3 washes with 1X PBS and the corresponding secondary antibody was incubated for one hour. Proteins were visualized by LI-COR (Odyssey Fc) measuring secondary antibody fluorescence or using Amersham ECL Prime Western Blotting Detection reagent (GE Lifesciences) for HRP-linked antibodies with an ImageQuant (LAS8000 Mini). Primary and secondary antibodies used in this study: 1:50 HIV anti-p24Gag [64] (Mouse), 1:3000 anti-HIV gp160/120 (Rabbit, ADP421; Centralized Facility for AIDS Reagents (CFAR), 1:5000 anti-HSP90 (Rabbit, GeneTex, GTX109753), 1:1000 anti-FLAG (DYKDDDDK, Rabbit, Cell Signaling, 14793), 1:2000 anti-β-actin (Mouse, Abcam; Ab6276), 1:5000 anti-ZAP (Rabbit, Abcam, ab154680), 1:1000 anti-GFP (Mouse. Roche 11814460001), 1:5000 anti-rabbit HRP (Cell Signaling Technology, 7074), 1:5000 anti-mouse HRP (Cell Signaling Technology, 7076), 1:5000 anti-mouse IRDye 680RD (LI-COR, 926–68070), 1:5000 anti-rabbit IRDye 800CW (LI-COR, 926–32211).

### TZM-bl infectivity assay

The TZM-bl indicator cell line was used to quantify the amount of infectious virus [65–67]. Briefly, cells were seeded in 24-well plates and infected by incubation with virus stocks. 48 hours post-infection, the cells were lysed and infectivity was measured by β-galactosidase expression using the Galacto-Star System following manufacturer’s instructions (Applied Biosystems). β-galactosidase activity was quantified as relative light units per second using a PerkinElmer Luminometer.

### Immunoprecipitation assays

For the UV crosslinking, immunoprecipitation and γ^32^P-ATP RNA labelling assay, KHNYN CRISPR HeLa cells stably expressing GFP or GFP-KHNYN were seeded in 10cm dishes 24 hours prior to UV crosslinking. Dynabeads protein G beads (ThermoFisher Scientific) were washed twice with lysis buffer (50 mM Tris–HCl, pH 7.4, 100 mM NaCl, 1% Igepal CA-630, 0.1% SDS, 0.5% sodium deoxycholate and protease inhibitor cocktail) and incubated with 5 μg anti-ZAP (Abcam, ab154680) antibody for 1 hour at 4°C. The cells were washed with cold 1X PBS, UV-crosslinked at 400 mJ/cm^2^ and collected by scraping. DNA was sheared with the Bioruptor Pico (Diagenode, B01060010) for 30 seconds on/off five times. The samples were then incubated with 4U of DNAse Turbo (Invitrogen AM2238) and 2,5 U/mL of RNAse I (Invitrogen, EN0602) for 5 minutes at 37°C with shaking (1100 rpm). Lysates were then centrifuged for 10 minutes at 15000xg and loaded on the antibody-conjugated beads overnight. The following day, the samples were washed twice with high-salt buffer (50 mM Tris–HCl, pH 7.4, 1M NaCl, 1% Igepal CA-630, 0.1% SDS, 0.5% sodium deoxycholate and protease inhibitor cocktail) and once with 1X FastAP buffer (10 mM Tris-HCl (pH 8.0 at 37 °C), 5mM MgCl2, 0.1M KCl, 0.02% Triton X-100 and 0.1 mg/mL BSA), prior to incubation with 0.5U of FastAP (Invitrogen, EF0654) for 40 minutes at 37°C and 1100 rpm shaking. Following FastAP digestion, the samples were washed twice with high-salt wash buffer and once with 1X PNK buffer (7 mM Tris-HCl pH 7.6, 1 mM MgCl2, 0.5 mM dithiothreitol), followed by RNA labelling with γ^32^P-ATP during 60 minutes at 37°C and 1100 rpm shaking. The radiolabeled beads were pelleted and suspended in 2X Laemmli buffer, incubated for 10 minutes at 70°C and then loaded into a 4-12% NuPAGE Bis-Tris gel (Invitrogen, NP0326BOX). After electrophoresis, the gel was transferred to a nitrocellulose membrane and visualized using a Typhoon TRIO.

For the UV crosslinking, immunoprecipitation and quantitative RT-PCR assays, control CRISPR, TRIM25 CRISPR and KHNYN CRISPR HeLa cells were plated in 6-well plates and transfected using TransIT-LT1 transfection reagent with 500ng of pcDNA3.1-GFP or pcDNA3.1-ZAP-L plus 500ng of pHIV-1WT or pHIV-1CpG. The media was changed 4-6 hours later. 48 hours post-transfection, the cells were washed with 1X PBS prior to ‘on-dish’ irradiation with 400mJ/cm2 using a UV Stralinker 2400. Cells were then pelleted and lysed with RIPA buffer containing 50mM Tris-HCl (pH 7.4), 150mM NaCl, 0.1% SDS, 0.5% sodium deoxycholate, 1% NP-40 and protease inhibitors (Roche), and then sonicated. Cleared lysates were immunoprecipitated overnight at 4°C with a rabbit anti-ZAP antibody (Abcam) and protein G beads (Thermo Fisher). Following three washes with RIPA buffer, beads were resuspended in 100μl of RIPA and boiled for 10 minutes to decouple protein/RNA complexes from the beads. Finally, input and pulldown samples were incubated with proteinase K (Thermo Fisher, 2mg/ml) for 1 hour at 37°C, and then boiled for 10 minutes to inactivate the enzyme. Samples were stored at −20°C for downstream RNA extraction and RT-qPCR analysis. RNA was isolated and purified from the lysates by first resuspending the input and pulldown samples in QIAzol (QIAGEN). The suspension was passed through QIAshredder columns (QIAGEN) for homogenization, and then transferred to phase lock gel tubes (VWR) prior to addition of chloroform (SIGMA). After manually shaking the tubes, samples were centrifuged full speed, for 15 minutes at 4°C. The aqueous phase was passed to a new tube, and isopropanol added. After 10 minutes at room temperature, tubes were centrifuged as before, and supernatants removed. RNA pellets were subsequently washed with 75% ethanol, and centrifuged at 7500 x g, for 5 minutes at 4°C. Following aspiration of the supernatants, the RNA pellets were left to dry and then resuspended in RNase-free water. Purified RNAs were reverse transcribed by random hexamer primers using a High-Capacity cDNA Reverse Transcription kit (Applied Biosystems), according to the manufacturer instructions. Of the final reaction, 5μl were used for quantitative PCR with primer/probe sets for human GAPDH (Applied Biosystems Cat# Hs99999905_m1) and HIV-1 genomic RNA (primers GGCCAGGGAATTTTCTTCAGA / TTGTCTCTTCCCCAAACCTGA (forward/reverse) and probe FAM-ACCAGAGCCAACAGCCCCACCAGA-TAMRA). Serial dilutions of HIV-1WT proviral DNA were used for standard curves to quantify HIV-1 RNA copies.

For the assays analyzing KHNYN-GFP co-immunoprecipitation with ZAP, HeLa cells stably expressing wild-type KHNYN-GFP wild-type or mutant versions were seeded in 6-well plates for 24 hours prior to immunoprecipitation. The cells were lysed on ice in lysis buffer (0.5% NP-40, 150 mM KCl, 10 mM HEPES pH 7.5, 3 mM MgCl2) supplemented with complete Protease inhibitor cocktail tablets (Sigma-Aldrich). The lysates were incubated on ice for 1 hour and centrifugated at 20,000 x g for 15 minutes at 4°C. 50 μl of the post-nuclear supernatant was saved as the input lysate and 450 μL was incubated with 5μg of anti-GFP antibody (Roche 11814460001) for 1 hour at 4°C. Protein G Dynabeads (Invitrogen) were then added and incubated overnight at 4°C with rotation. The lysates were washed four times with wash buffer (0.05% NP-40, 150 mM KCl, 10 mM HEPES pH 7.5, 3 mM MgCl_2_) before the bound proteins were eluted with 2X Laemmli buffer and boiled for 10 minutes. Protein expression was analyzed by western blotting as described above.

### Microscopy

HeLa cells were seeded on 24-well plates on coverslips pre-treated with poly-lysine. KHNYN CRISPR HeLa cells were transfected with 250ng of CRISPR-resistant KHNYN-FLAG, ΔKH-KHNYN-FLAG or GDDG-KHNYN-FLAG. 24 hours post-transfection, the cells were fixed with 4% paraformaldehyde for 20 minutes at room temperature, washed once with 1X PBS and once in 10mM glycine. Cells were then permeabilized for 15 minutes in 1% BSA and 0.1% Triton-X in PBS. HeLa cells stably expressing wild-type KHNYN-GFP or versions with specific mutations were seeded in pre-treated 24-well plates 24 hours prior to immunostaining and were fixed and permeabilized as above. Mouse anti-FLAG (1:500), rabbit anti-ZAP (1:500) or rabbit anti-TRIM25 (1:500) antibodies were diluted in 1X PBS/0.01% Triton-X and the cells were stained for 1 hour at room temperature. The cells were then washed three times in PBS/0.01% Triton-X and incubated with Alexa Fluor 594 anti-mouse or Alexa Fluor 647 anti-rabbit (Molecular Probes, 1:500 in 1X PBS/0.01% Triton-X) for 45 minutes in the dark. Finally, the coverslips were washed three times with 1X PBS/0.01% Triton-X and then mounted on slides using Prolong Diamond Antifade Mountant with DAPI (Invitrogen). Imaging was performed on a Nikon Eclipse Ti Inverted Microscope, equipped with a Yokogawa CSU/X1-spinning disk unit, under 100x objective and laser wavelengths of 405 nm, 488 nm, 561 nm, and 640 nm. Image processing and co-localization analysis was performed with Image J (Fiji) software.

For the Leptomycin B treatment experiments, HeLa cells stably expressing KHNYN-GFP wild-type or mutants were seeded in pre-treated 24-well plates 24 hours prior to 4 hour treatment with 50nM of Leptomycin B or DMSO at 37°C. After treatment, the cells were fixed and immunostained as described above.

### KHNYN domain prediction, KH domain alignment, NES identification and phylogenetic analysis of KHNYN and N4BP1

Phyre2 was used on the full-length KHNYN sequence (NP_056114) using the intensive modelling mode [37]. KH domains were aligned using MUSCLE [68] implemented in the DNA STAR suite of programs. The NES was identified using the Wregex tool with the NES/CRM1 motif and the relaxed configuration [59]. Amino acid sequences for KHNYN and N4BP1 were obtained from NCBI Gene, checked manually to ensure they were full-length sequences and aligned using ClustalOmega. The resulting alignment file was used to infer a maximum likelihood tree in the DIVEIN web server [69] using the LG substitution model, and the N4BP1-like sequences from the Californian sea hare (*Aplysia californica*), and the Crown-of-thorns Starfish (*Acanthaster planci*) as outgroups. The resulting tree was visually presented and annotated using the interactive Tree of life (iTol) [70].

### Statistical analysis

Statistical significance was determined using unpaired two-tailed *t*-tests in GraphPad. Data are represented as mean ± standard deviation and significance was ascribed to p values p < 0.05.

## ACKNOWLEDGEMENTS

We thank members of the Neil and Swanson laboratories as well as Michael Malim for helpful discussions. The following reagents were obtained through the NIH AIDS Research and Reference Reagent Program, Division of AIDS, NIAID, NIH: TZM-bl from Dr. John C. Kappes, Dr. Xiaoyun Wu and Tranzyme Inc; HIV-1 p24 Hybridoma (183-H12-5C) from Dr. Bruce Chesebro. The Antiserum to HIV-1 gp120 #20 (ARP421) was obtained from the NIBSC Centre for AIDS Reagents. These studies were funded by a Medical Research Council grant MR/S000844/1 to SJDN and CMS, a Deutsche Forschungsgemeinschaft (German Research Foundation) fellowship to DK (Project number: KM 5/1-1), a Wellcome Trust Senior Research Fellowship (WT098049AIA) to SJDN, a Royal Society/Wellcome Trust Sir Henry Dale Fellowship (206200/Z/17/Z) to CO and the Francis Crick Institute, which receives its core funding from Cancer Research UK (FC001178), the UK Medical Research Council (FC001178) and the Wellcome Trust (FC001178). MF is supported by the UK Medical Research Council (MR/R50225X/1) and is a King’s College London member of the MRC Doctoral Training Partnership in Biomedical Sciences. This work was supported by the Department of Health via a National Institute for Health Research Comprehensive Biomedical Research Centre award to Guy’s and St. Thomas’ NHS Foundation Trust in partnership with King’s College London and King’s College Hospital NHS Foundation Trust.

## COMPETING INTERESTS

The authors declare no competing interests.

## SUPPLEMENTAL FIGURE LEGENDS

**Figure S1. Deletion of the extended di-KH domain reduces KHNYN antiviral activity on viral gene expression.** Representative western blots for HIV-1 Gag and Env expression in cell lysates as well as virion production for the experiments shown in Figure 2B.

**Figure S2: Mutation of the GxxG motif in KH2 does not reduce KHNYN antiviral activity on viral gene expression.** Representative western blots for HIV-1 Gag and Env expression in cell lysates as well as virion production for the experiments shown in Figure 3B.

**Figure S3: Characterization of KHNYN-GFP cell lines and KHNYN is not required for overexpressed ZAP to bind HIV-1 RNA. (A)** Expression of proteins in HeLa cells that have been reported to regulate ZAP RNA degradation. Data is from reference [51]. **(B)** Control CRISPR, ZAP CRISPR, KHNYN CRISPR or KHNYN CRISPR + KHNYN-GFP cells were infected with VSV-G pseudotyped HIV-1_WT_ or HIV-1_CpG_ with an MOI of 1. 48 hours post-infection, cell supernatant was harvested and infectious virus production was measured in TZM-bl cells. Each bar shows the average value of three independent experiments normalized to the value obtained for wild-type KHNYN co-transfected with pHIV-1_WT_. *p < 0.05 as determined by a two-tailed unpaired t-test comparing HIV-1_CpG_ in each cell line to the control CRISPR cell line. **(C)** Western blot for KHNYN in control CRISPR, KHNYN CRISPR and KHNYN CRISPR + KHNYN-GFP cells. **(D)** Quantification of HIV-1 genomic RNA (gRNA) bound to ZAP after crosslinking and immunoprecipitation in control CRISPR cells, TRIM25 CRISPR cells or KHNYN CRISPR cells co-transfected with pZAP-L and either pHIV-1_WT_ or pHIV-1_CpG_. Each bar shows the average value of three independent experiments normalized to the value obtained for pHIV-1_WT_ in each cell line. *p < 0.05 as determined by a two-tailed unpaired t-test comparing HIV-1_CpG_ to HIV-1_WT_ in each cell line.

**Figure S4. The C-terminal CUBAN domain is essential for KHNYN antiviral activity on viral gene expression. (A)** Representative western blots for Gag and Env in cell lysates as well as virion production for the experiments shown in Figure 4B. **(B)** Representative western blots for Gag and Env in cell lysates as well as virion production for the experiments shown in Figure 4C.

**Figure S5. The nuclear export signal at the C-terminus of the KHNYN CUBAN domain is required for antiviral activity on viral gene expression. (A)** Representative western blots for Gag and Env in cell lysates as well as virion production for the experiments shown in Figure 6C. **(B)** Confocal microscopy images for wild-type KHNYN-GFP, KHNYNΔCUBAN-GFP, KHNYN-ΔNES and KHNYN-NESmut-GFP (green) co-staining with endogenous TRIM25 (magenta), scale bar is 10μm.

**Figure S6. The nuclear export signal in KHNYN is conserved in mammals, reptiles and amphibians.** Maximum likelihood phylogenetic tree of KHNYN and N4BP1 amino acid sequences. Representative sequences from bony fish (orange), amphibians (green), reptiles (yellow), birds (light purple) and mammals (light blue) were aligned and maximum likelihood phylogeny was inferred using the LG substitution model DIVEIN. The crown-of-thorns starfish and Californian sea hare were used as outgroups to root the tree. (+) all five positions of the PKI-type NES in the C-terminus of KHNYN are present in the lineage. (-) positions 3 and 4 in the NES are not present in the lineage.

## Notes

### Competing Interest Statement

The authors have declared no competing interest.

## REFERENCES

1. Ficarelli M, Neil SJD, Swanson CM. Targeted Restriction of Viral Gene Expression and Replication by the ZAP Antiviral System. Annu Rev Virol. 2021;8(1):265–83. Epub 2021/06/16. doi: 10.1146/annurev-virology-091919-104213. PubMed PMID: 34129371.

2. Hur S. Double-Stranded RNA Sensors and Modulators in Innate Immunity. Annual review of immunology. 2019;37:349–75. Epub 2019/01/24. doi: 10.1146/annurev-immunol-042718-041356. PubMed PMID: 30673536; PubMed Central PMCID: PMCPMC7136661.

3. Meagher JL, Takata M, Goncalves-Carneiro D, Keane SC, Rebendenne A, Ong H, et al. Structure of the zinc-finger antiviral protein in complex with RNA reveals a mechanism for selective targeting of CG-rich viral sequences. Proc Natl Acad Sci U S A. 2019;116(48):24303–9. Epub 2019/11/14. doi: 10.1073/pnas.1913232116. PubMed PMID: 31719195; PubMed Central PMCID: PMCPMC6883784.

4. Luo X, Wang X, Gao Y, Zhu J, Liu S, Gao G, et al. Molecular Mechanism of RNA Recognition by Zinc-Finger Antiviral Protein. Cell reports. 2020;30(1):46–52 e4. Epub 2020/01/09. doi: 10.1016/j.celrep.2019.11.116. PubMed PMID: 31914396.

5. Greenbaum BD, Rabadan R, Levine AJ. Patterns of oligonucleotide sequences in viral and host cell RNA identify mediators of the host innate immune system. PLoS One. 2009;4(6):e5969. Epub 2009/06/19. doi: 10.1371/journal.pone.0005969. PubMed PMID: 19536338; PubMed Central PMCID: PMCPMC2694999.

6. Tulloch F, Atkinson NJ, Evans DJ, Ryan MD, Simmonds P. RNA virus attenuation by codon pair deoptimisation is an artefact of increases in CpG/UpA dinucleotide frequencies. Elife. 2014;3:e04531. Epub 2014/12/10. doi: 10.7554/eLife.04531. PubMed PMID: 25490153; PubMed Central PMCID: PMCPMC4383024.

7. Guo X, Ma J, Sun J, Gao G. The zinc-finger antiviral protein recruits the RNA processing exosome to degrade the target mRNA. Proc Natl Acad Sci U S A. 2007;104(1):151–6. Epub 2006/12/23. doi: 10.1073/pnas.0607063104. PubMed PMID: 17185417; PubMed Central PMCID: PMCPMC1765426.

8. Zhu Y, Chen G, Lv F, Wang X, Ji X, Xu Y, et al. Zinc-finger antiviral protein inhibits HIV-1 infection by selectively targeting multiply spliced viral mRNAs for degradation. Proc Natl Acad Sci U S A. 2011;108(38):15834–9. Epub 2011/08/31. doi: 10.1073/pnas.1101676108. PubMed PMID: 21876179; PubMed Central PMCID: PMCPMC3179061.

9. Li MM, Lau Z, Cheung P, Aguilar EG, Schneider WM, Bozzacco L, et al. TRIM25 Enhances the Antiviral Action of Zinc-Finger Antiviral Protein (ZAP). PLoS Pathog. 2017;13(1):e1006145. Epub 2017/01/07. doi: 10.1371/journal.ppat.1006145. PubMed PMID: 28060952; PubMed Central PMCID: PMCPMC5245905.

10. Zheng X, Wang X, Tu F, Wang Q, Fan Z, Gao G. TRIM25 Is Required for the Antiviral Activity of Zinc Finger Antiviral Protein. J Virol. 2017;91(9):e00088–17. Epub 2017/02/17. doi: 10.1128/JVI.00088-17. PubMed PMID: 28202764; PubMed Central PMCID: PMCPMC5391446.

11. Zhu Y, Wang X, Goff SP, Gao G. Translational repression precedes and is required for ZAP-mediated mRNA decay. EMBO J. 2012;31(21):4236–46. Epub 2012/10/02. doi: 10.1038/emboj.2012.271. PubMed PMID: 23023399; PubMed Central PMCID: PMCPMC3492732.

12. Ficarelli M, Wilson H, Pedro Galao R, Mazzon M, Antzin-Anduetza I, Marsh M, et al. KHNYN is essential for the zinc finger antiviral protein (ZAP) to restrict HIV-1 containing clustered CpG dinucleotides. Elife. 2019;8:e46767. Epub 2019/07/10. doi: 10.7554/eLife.46767. PubMed PMID: 31284899; PubMed Central PMCID: PMCPMC6615859.

13. Anantharaman V, Aravind L. The NYN domains: novel predicted RNAses with a PIN domain-like fold. RNA Biol. 2006;3(1):18–27. Epub 2006/11/23. doi: 10.4161/rna.3.1.2548. PubMed PMID: 17114934.

14. Matelska D, Steczkiewicz K, Ginalski K. Comprehensive classification of the PIN domain-like superfamily. Nucleic Acids Res. 2017;45(12):6995–7020. Epub 2017/06/03. doi: 10.1093/nar/gkx494. PubMed PMID: 28575517; PubMed Central PMCID: PMCPMC5499597.

15. Castagnoli L, Mandaliti W, Nepravishta R, Valentini E, Mattioni A, Procopio R, et al. Selectivity of the CUBAN domain in the recognition of ubiquitin and NEDD8. FEBS J. 2019;286(4):653–77. Epub 2019/01/20. doi: 10.1111/febs.14752. PubMed PMID: 30659753.

16. Marco A, Marin I. CGIN1: a retroviral contribution to mammalian genomes. Mol Biol Evol. 2009;26(10):2167–70. Epub 2009/06/30. doi: 10.1093/molbev/msp127. PubMed PMID: 19561090.

17. Shaw AE, Hughes J, Gu Q, Behdenna A, Singer JB, Dennis T, et al. Fundamental properties of the mammalian innate immune system revealed by multispecies comparison of type I interferon responses. PLoS Biol. 2017;15(12):e2004086. Epub 2017/12/19. doi: 10.1371/journal.pbio.2004086. PubMed PMID: 29253856; PubMed Central PMCID: PMCPMC5747502.

18. Nepravishta R, Ferrentino F, Mandaliti W, Mattioni A, Weber J, Polo S, et al. CoCUN, a Novel Ubiquitin Binding Domain Identified in N4BP1. Biomolecules. 2019;9(7). Epub 2019/07/20. doi: 10.3390/biom9070284. PubMed PMID: 31319543; PubMed Central PMCID: PMCPMC6681339.

19. Murillas R, Simms KS, Hatakeyama S, Weissman AM, Kuehn MR. Identification of developmentally expressed proteins that functionally interact with Nedd4 ubiquitin ligase. J Biol Chem. 2002;277(4):2897–907. Epub 2001/11/22. doi: 10.1074/jbc.M110047200. PubMed PMID: 11717310.

20. Sharma P, Murillas R, Zhang H, Kuehn MR. N4BP1 is a newly identified nucleolar protein that undergoes SUMO-regulated polyubiquitylation and proteasomal turnover at promyelocytic leukemia nuclear bodies. J Cell Sci. 2010;123(Pt 8):1227–34. Epub 2010/03/18. doi: 10.1242/jcs.060160. PubMed PMID: 20233849; PubMed Central PMCID: PMCPMC2848111.

21. Oberst A, Malatesta M, Aqeilan RI, Rossi M, Salomoni P, Murillas R, et al. The Nedd4-binding partner 1 (N4BP1) protein is an inhibitor of the E3 ligase Itch. Proc Natl Acad Sci U S A. 2007;104(27):11280–5. Epub 2007/06/27. doi: 10.1073/pnas.0701773104. PubMed PMID: 17592138; PubMed Central PMCID: PMCPMC2040890.

22. Yamasoba D, Sato K, Ichinose T, Imamura T, Koepke L, Joas S, et al. N4BP1 restricts HIV-1 and its inactivation by MALT1 promotes viral reactivation. Nat Microbiol. 2019;4(9):1532–44. Epub 2019/05/28. doi: 10.1038/s41564-019-0460-3. PubMed PMID: 31133753.

23. Gitlin AD, Heger K, Schubert AF, Reja R, Yan D, Pham VC, et al. Integration of innate immune signalling by caspase-8 cleavage of N4BP1. Nature. 2020;587(7833):275–80. Epub 2020/09/25. doi: 10.1038/s41586-020-2796-5. PubMed PMID: 32971525.

24. Shi H, Sun L, Wang Y, Liu A, Zhan X, Li X, et al. N4BP1 negatively regulates NF-kappaB by binding and inhibiting NEMO oligomerization. Nature communications. 2021;12(1):1379. Epub 2021/03/04. doi: 10.1038/s41467-021-21711-5. PubMed PMID: 33654074; PubMed Central PMCID: PMCPMC7925594.

25. OhAinle M, Helms L, Vermeire J, Roesch F, Humes D, Basom R, et al. A virus-packageable CRISPR screen identifies host factors mediating interferon inhibition of HIV. Elife. 2018;7:e39823. Epub 2018/12/07. doi: 10.7554/eLife.39823. PubMed PMID: 30520725; PubMed Central PMCID: PMCPMC6286125.

26. Kypr J, Mrazek J, Reich J. Nucleotide composition bias and CpG dinucleotide content in the genomes of HIV and HTLV 1/2. Biochim Biophys Acta. 1989;1009(3):280–2. Epub 1989/12/22. doi: 10.1016/0167-4781(89)90114-0. PubMed PMID: 2597678.

27. Shpaer EG, Mullins JI. Selection against CpG dinucleotides in lentiviral genes: a possible role of methylation in regulation of viral expression. Nucleic Acids Res. 1990;18(19):5793–7. Epub 1990/10/11. doi: 10.1093/nar/18.19.5793. PubMed PMID: 2170945; PubMed Central PMCID: PMCPMC332316.

28. Takata MA, Goncalves-Carneiro D, Zang TM, Soll SJ, York A, Blanco-Melo D, et al. CG dinucleotide suppression enables antiviral defence targeting non-self RNA. Nature. 2017;550(7674):124–7. Epub 2017/09/28. doi: 10.1038/nature24039. PubMed PMID: 28953888; PubMed Central PMCID: PMCPMC6592701.

29. Ficarelli M, Antzin-Anduetza I, Hugh-White R, Firth AE, Sertkaya H, Wilson H, et al. CpG Dinucleotides Inhibit HIV-1 Replication through Zinc Finger Antiviral Protein (ZAP)-Dependent and -Independent Mechanisms. J Virol. 2020;94(6):e01337–19. Epub 2019/11/22. doi: 10.1128/JVI.01337-19. PubMed PMID: 31748389; PubMed Central PMCID: PMCPMC7158733.

30. Kmiec D, Nchioua R, Sherrill-Mix S, Sturzel CM, Heusinger E, Braun E, et al. CpG Frequency in the 5’ Third of the env Gene Determines Sensitivity of Primary HIV-1 Strains to the Zinc-Finger Antiviral Protein. mBio. 2020;11(1):e02903–19. Epub 2020/01/16. doi: 10.1128/mBio.02903-19. PubMed PMID: 31937644; PubMed Central PMCID: PMCPMC6960287.

31. Gao G, Guo X, Goff SP. Inhibition of retroviral RNA production by ZAP, a CCCH-type zinc finger protein. Science. 2002;297(5587):1703–6. Epub 2002/09/07. doi: 10.1126/science.1074276. PubMed PMID: 12215647.

32. Kerns JA, Emerman M, Malik HS. Positive selection and increased antiviral activity associated with the PARP-containing isoform of human zinc-finger antiviral protein. PLoS genetics. 2008;4(1):e21. Epub 2008/01/30. doi: 10.1371/journal.pgen.0040021. PubMed PMID: 18225958; PubMed Central PMCID: PMCPMC2213710.

33. Charron G, Li MM, MacDonald MR, Hang HC. Prenylome profiling reveals S-farnesylation is crucial for membrane targeting and antiviral activity of ZAP long-isoform. Proc Natl Acad Sci U S A. 2013;110(27):11085–90. Epub 2013/06/19. doi: 10.1073/pnas.1302564110. PubMed PMID: 23776219; PubMed Central PMCID: PMCPMC3703996.

34. Schwerk J, Soveg FW, Ryan AP, Thomas KR, Hatfield LD, Ozarkar S, et al. RNA-binding protein isoforms ZAP-S and ZAP-L have distinct antiviral and immune resolution functions. Nat Immunol. 2019;20(12):1610–20. Epub 2019/11/20. doi: 10.1038/s41590-019-0527-6. PubMed PMID: 31740798; PubMed Central PMCID: PMCPMC7240801.

35. Kmiec D, Lista MJ, Ficarelli M, Swanson CM, Neil SJD. S-farnesylation is essential for antiviral activity of the long ZAP isoform against RNA viruses with diverse replication strategies. PLoS Pathog. 2021;17(10):e1009726. Epub 2021/10/26. doi: 10.1371/journal.ppat.1009726. PubMed PMID: 34695163; PubMed Central PMCID: PMCPMC8568172.

36. Go CD, Knight JDR, Rajasekharan A, Rathod B, Hesketh GG, Abe KT, et al. A proximity-dependent biotinylation map of a human cell. Nature. 2021;595(7865):120–4. Epub 2021/06/04. doi: 10.1038/s41586-021-03592-2. PubMed PMID: 34079125.

37. Kelley LA, Mezulis S, Yates CM, Wass MN, Sternberg MJ. The Phyre2 web portal for protein modeling, prediction and analysis. Nat Protoc. 2015;10(6):845–58. Epub 2015/05/08. doi: 10.1038/nprot.2015.053. PubMed PMID: 25950237; PubMed Central PMCID: PMCPMC5298202.

38. Jumper J, Evans R, Pritzel A, Green T, Figurnov M, Ronneberger O, et al. Highly accurate protein structure prediction with AlphaFold. Nature. 2021;596(7873):583–9. Epub 2021/07/16. doi: 10.1038/s41586-021-03819-2. PubMed PMID: 34265844; PubMed Central PMCID: PMCPMC8371605.

39. Nicastro G, Taylor IA, Ramos A. KH-RNA interactions: back in the groove. Curr Opin Struct Biol. 2015;30:63–70. Epub 2015/01/28. doi: 10.1016/j.sbi.2015.01.002. PubMed PMID: 25625331.

40. Braddock DT, Louis JM, Baber JL, Levens D, Clore GM. Structure and dynamics of KH domains from FBP bound to single-stranded DNA. Nature. 2002;415(6875):1051–6. Epub 2002/03/05. doi: 10.1038/4151051a. PubMed PMID: 11875576.

41. Nicastro G, Candel AM, Uhl M, Oregioni A, Hollingworth D, Backofen R, et al. Mechanism of beta-actin mRNA Recognition by ZBP1. Cell reports. 2017;18(5):1187–99. Epub 2017/02/02. doi: 10.1016/j.celrep.2016.12.091. PubMed PMID: 28147274; PubMed Central PMCID: PMCPMC5300891.

42. Diaz-Moreno I, Hollingworth D, Kelly G, Martin S, Garcia-Mayoral M, Briata P, et al. Orientation of the central domains of KSRP and its implications for the interaction with the RNA targets. Nucleic Acids Res. 2010;38(15):5193–205. Epub 2010/04/14. doi: 10.1093/nar/gkq216. PubMed PMID: 20385598; PubMed Central PMCID: PMCPMC2926597.

43. Chao JA, Patskovsky Y, Patel V, Levy M, Almo SC, Singer RH. ZBP1 recognition of beta-actin zipcode induces RNA looping. Genes Dev. 2010;24(2):148–58. Epub 2010/01/19. doi: 10.1101/gad.1862910. PubMed PMID: 20080952; PubMed Central PMCID: PMCPMC2807350.

44. Dagil R, Ball NJ, Ogrodowicz RW, Hobor F, Purkiss AG, Kelly G, et al. IMP1 KH1 and KH2 domains create a structural platform with unique RNA recognition and re-modelling properties. Nucleic Acids Res. 2019;47(8):4334–48. Epub 2019/03/14. doi: 10.1093/nar/gkz136. PubMed PMID: 30864660; PubMed Central PMCID: PMCPMC6486635.

45. Gough J, Karplus K, Hughey R, Chothia C. Assignment of homology to genome sequences using a library of hidden Markov models that represent all proteins of known structure. J Mol Biol. 2001;313(4):903–19. Epub 2001/11/08. doi: 10.1006/jmbi.2001.5080. PubMed PMID: 11697912.

46. Howe KL, Achuthan P, Allen J, Allen J, Alvarez-Jarreta J, Amode MR, et al. Ensembl 2021. Nucleic Acids Res. 2021;49(D1):D884–D91. Epub 2020/11/03. doi: 10.1093/nar/gkaa942. PubMed PMID: 33137190; PubMed Central PMCID: PMCPMC7778975.

47. Oddone A, Lorentzen E, Basquin J, Gasch A, Rybin V, Conti E, et al. Structural and biochemical characterization of the yeast exosome component Rrp40. EMBO Rep. 2007;8(1):63–9. Epub 2006/12/13. doi: 10.1038/sj.embor.7400856. PubMed PMID: 17159918; PubMed Central PMCID: PMCPMC1796750.

48. Nakel K, Hartung SA, Bonneau F, Eckmann CR, Conti E. Four KH domains of the C. elegans Bicaudal-C ortholog GLD-3 form a globular structural platform. RNA. 2010;16(11):2058–67. Epub 2010/09/09. doi: 10.1261/rna.2315010. PubMed PMID: 20823118; PubMed Central PMCID: PMCPMC2957046.

49. Hollingworth D, Candel AM, Nicastro G, Martin SR, Briata P, Gherzi R, et al. KH domains with impaired nucleic acid binding as a tool for functional analysis. Nucleic Acids Res. 2012;40(14):6873–86. Epub 2012/05/02. doi: 10.1093/nar/gks368. PubMed PMID: 22547390; PubMed Central PMCID: PMCPMC3413153.

50. Gebauer F, Schwarzl T, Valcarcel J, Hentze MW. RNA-binding proteins in human genetic disease. Nat Rev Genet. 2021;22(3):185–98. Epub 2020/11/26. doi: 10.1038/s41576-020-00302-y. PubMed PMID: 33235359.

51. Nagaraj N, Wisniewski JR, Geiger T, Cox J, Kircher M, Kelso J, et al. Deep proteome and transcriptome mapping of a human cancer cell line. Molecular systems biology. 2011;7:548. Epub 2011/11/10. doi: 10.1038/msb.2011.81. PubMed PMID: 22068331; PubMed Central PMCID: PMCPMC3261714.

52. Ryman KD, Meier KC, Nangle EM, Ragsdale SL, Korneeva NL, Rhoads RE, et al. Sindbis virus translation is inhibited by a PKR/RNase L-independent effector induced by alpha/beta interferon priming of dendritic cells. J Virol. 2005;79(3):1487–99. Epub 2005/01/15. doi: 10.1128/JVI.79.3.1487-1499.2005. PubMed PMID: 15650175; PubMed Central PMCID: PMCPMC544143.

53. Nchioua R, Kmiec D, Muller JA, Conzelmann C, Gross R, Swanson CM, et al. SARS-CoV-2 Is Restricted by Zinc Finger Antiviral Protein despite Preadaptation to the Low-CpG Environment in Humans. mBio. 2020;11(5):e01930–20. Epub 2020/10/18. doi: 10.1128/mBio.01930-20. PubMed PMID: 33067384; PubMed Central PMCID: PMCPMC7569149.

54. Guttler T, Gorlich D. Ran-dependent nuclear export mediators: a structural perspective. EMBO J. 2011;30(17):3457–74. Epub 2011/09/01. doi: 10.1038/emboj.2011.287. PubMed PMID: 21878989; PubMed Central PMCID: PMCPMC3181476.

55. Kirli K, Karaca S, Dehne HJ, Samwer M, Pan KT, Lenz C, et al. A deep proteomics perspective on CRM1-mediated nuclear export and nucleocytoplasmic partitioning. Elife. 2015;4. Epub 2015/12/18. doi: 10.7554/eLife.11466. PubMed PMID: 26673895; PubMed Central PMCID: PMCPMC4764573.

56. Kudo N, Wolff B, Sekimoto T, Schreiner EP, Yoneda Y, Yanagida M, et al. Leptomycin B inhibition of signal-mediated nuclear export by direct binding to CRM1. Exp Cell Res. 1998;242(2):540–7. Epub 1998/07/31. doi: 10.1006/excr.1998.4136. PubMed PMID: 9683540.

57. Liu L, Chen G, Ji X, Gao G. ZAP is a CRM1-dependent nucleocytoplasmic shuttling protein. Biochem Biophys Res Commun. 2004;321(3):517–23. Epub 2004/09/11. doi: 10.1016/j.bbrc.2004.06.174. PubMed PMID: 15358138.

58. Guttler T, Madl T, Neumann P, Deichsel D, Corsini L, Monecke T, et al. NES consensus redefined by structures of PKI-type and Rev-type nuclear export signals bound to CRM1. Nat Struct Mol Biol. 2010;17(11):1367–76. Epub 2010/10/26. doi: 10.1038/nsmb.1931. PubMed PMID: 20972448.

59. Prieto G, Fullaondo A, Rodriguez JA. Prediction of nuclear export signals using weighted regular expressions (Wregex). Bioinformatics. 2014;30(9):1220–7. Epub 2014/01/15. doi: 10.1093/bioinformatics/btu016. PubMed PMID: 24413524.

60. Goncalves-Carneiro D, Takata MA, Ong H, Shilton A, Bieniasz PD. Origin and evolution of the zinc finger antiviral protein. PLoS Pathog. 2021;17(4):e1009545. Epub 2021/04/27. doi: 10.1371/journal.ppat.1009545. PubMed PMID: 33901262.

61. Antzin-Anduetza I, Mahiet C, Granger LA, Odendall C, Swanson CM. Increasing the CpG dinucleotide abundance in the HIV-1 genomic RNA inhibits viral replication. Retrovirology. 2017;14(1):49. Epub 2017/11/11. doi: 10.1186/s12977-017-0374-1. PubMed PMID: 29121951; PubMed Central PMCID: PMCPMC5679385.

62. Pear WS, Miller JP, Xu L, Pui JC, Soffer B, Quackenbush RC, et al. Efficient and rapid induction of a chronic myelogenous leukemia-like myeloproliferative disease in mice receiving P210 bcr/abl-transduced bone marrow. Blood. 1998;92(10):3780–92. Epub 1998/11/10. PubMed PMID: 9808572.

63. Fouchier RA, Meyer BE, Simon JH, Fischer U, Malim MH. HIV-1 infection of non-dividing cells: evidence that the amino-terminal basic region of the viral matrix protein is important for Gag processing but not for post-entry nuclear import. EMBO J. 1997;16(15):4531–9. Epub 1997/08/01. doi: 10.1093/emboj/16.15.4531. PubMed PMID: 9303297; PubMed Central PMCID: PMCPMC1170079.

64. Chesebro B, Wehrly K, Nishio J, Perryman S. Macrophage-tropic human immunodeficiency virus isolates from different patients exhibit unusual V3 envelope sequence homogeneity in comparison with T-cell-tropic isolates: definition of critical amino acids involved in cell tropism. J Virol. 1992;66(11):6547–54. Epub 1992/11/01. doi: 10.1128/JVI.66.11.6547-6554.1992. PubMed PMID: 1404602; PubMed Central PMCID: PMCPMC240149.

65. Derdeyn CA, Decker JM, Sfakianos JN, Wu X, O’Brien WA, Ratner L, et al. Sensitivity of human immunodeficiency virus type 1 to the fusion inhibitor T-20 is modulated by coreceptor specificity defined by the V3 loop of gp120. J Virol. 2000;74(18):8358–67. Epub 2000/08/23. doi: 10.1128/jvi.74.18.8358-8367.2000. PubMed PMID: 10954535; PubMed Central PMCID: PMCPMC116346.

66. Wei X, Decker JM, Liu H, Zhang Z, Arani RB, Kilby JM, et al. Emergence of resistant human immunodeficiency virus type 1 in patients receiving fusion inhibitor (T-20) monotherapy. Antimicrob Agents Chemother. 2002;46(6):1896–905. Epub 2002/05/23. doi: 10.1128/aac.46.6.1896-1905.2002. PubMed PMID: 12019106; PubMed Central PMCID: PMCPMC127242.

67. Platt EJ, Wehrly K, Kuhmann SE, Chesebro B, Kabat D. Effects of CCR5 and CD4 cell surface concentrations on infections by macrophagetropic isolates of human immunodeficiency virus type 1. J Virol. 1998;72(4):2855–64. Epub 1998/04/03. doi: 10.1128/JVI.72.4.2855-2864.1998. PubMed PMID: 9525605; PubMed Central PMCID: PMCPMC109730.

68. Edgar RC. MUSCLE: multiple sequence alignment with high accuracy and high throughput. Nucleic Acids Res. 2004;32(5):1792–7. Epub 2004/03/23. doi: 10.1093/nar/gkh340. PubMed PMID: 15034147; PubMed Central PMCID: PMCPMC390337.

69. Deng W, Maust BS, Nickle DC, Learn GH, Liu Y, Heath L, et al. DIVEIN: a web server to analyze phylogenies, sequence divergence, diversity, and informative sites. Biotechniques. 2010;48(5):405–8. Epub 2010/06/24. doi: 10.2144/000113370. PubMed PMID: 20569214; PubMed Central PMCID: PMCPMC3133969.

70. Letunic I, Bork P. Interactive Tree Of Life (iTOL) v5: an online tool for phylogenetic tree display and annotation. Nucleic Acids Res. 2021;49(W1):W293–W6. Epub 2021/04/23. doi: 10.1093/nar/gkab301. PubMed PMID: 33885785; PubMed Central PMCID: PMCPMC8265157.

